# Selection in parental species predicts hybrid evolution

**DOI:** 10.1101/283689

**Authors:** Anna Runemark, Richard I Bailey, Lena Bache-Mathiesen, Glenn-Peter Saetre

## Abstract

While hybridization is recognized as important in evolution, its contribution to adaptation and diversification remains poorly understood. Using genomically diverged island populations of the homoploid hybrid Italian sparrow, we test predictions for phenotypic trait values and evolvability based on patterns of parental species divergence in four plumage color traits. Fixed major QTL in species differences, favoured by strong selection, are expected to lead to hybrids with higher evolvability than the parent species. We find associations between parental divergence and trait evolution in Italian sparrows. Rump color shows little evidence of major QTL, and hybrid evolution closely matches parental variability. Back and crown plumage, however, show evidence of major QTL in species differences. For these traits, Italian sparrow phenotypes are biased towards axes of high parental differentiation and show greater phenotypic novelty along axes of low current parental evolvability, as predicted when major QTL are involved in species differences. Crown color has consistently evolved back towards one parent, while back color varies among islands. We also find among-island diversification within the Italian sparrow. Hence, hybridization of the same parent species can generate different phenotypes. In conclusion, we find that parental phenotypic divergence patterns can be useful in predicting hybrid evolutionary potential.

## Introduction

Understanding adaptation and diversification is central to evolutionary biology. The ability of populations to adapt to local conditions and of separate populations to differentiate and eventually allow for the formation of new species is influenced by the amount of heritable genetic variation present within those populations. Although hybridization is increasingly recognized as important in evolution, we still know little about how variation arising through hybridization contributes to divergence and adaptation. Here we investigate evolution within a hybrid species to address whether hybrid evolution and diversification can be predicted by patterns of phenotypic divergence between the parent species.

Population differentiation, ecological adaptation, and sexual selection all depend on additive genetic variation in traits combined with the strength of constraints among them. Different traits may affect each other’s evolution through being genetically correlated, for example due to pleiotropic loci, or through being under correlated or antagonistic selection (Guillaume and Whitlock 2007; Hansen and Houle 2008; Kirkpatrick 2009; Walsh and Blows 2009). Along with the magnitude of additive genetic variation, these constraints will influence the response to selection. In general, evolutionary change is expected to occur primarily along the axes with highest within-population additive genetic variance or ‘evolvability’ (Hansen and Houle 2008; Chenoweth et al. 2010; see Glossary), referred to as “lines of least resistance” (Schluter 1996; Glossary). Interestingly, hybridization may alter these lines as it can alter phenotypic trait variances and covariances (Rieseberg et al. 1999; Bailey et al. 2013; Selz et al. 2013; Lucek et al. 2016).

At the species level, hybridization is a common phenomenon (Mallet 2005; Abbott et al. 2013) and data from novel sequencing technologies show that many genomes are admixed (Pennisi 2016). We now know that hybridization can allow the transfer of adaptive genetic variants across species boundaries (Song et al. 2011; Heliconius Sequencing Consortium 2012), generate new species (Rieseberg 1997; Gompert et al. 2006), and spur adaptive radiations (Seehausen 2004; 2013; Meier et al. 2017). However, little is known about how hybridization-derived variation contributes to the ability to adapt and evolve (Bailey et al. 2013), but see Selz et al. (2013) and Lucek et al. (2016). Insights into the evolutionary potential of hybrid species would increase our understanding of the impact of hybridization on evolution, adaptation and diversity.

The phenotypic variation derived from hybridization is expected to differ from that derived from mutation. For instance, introgressed variants have been tested by evolution, albeit in a different genomic background. Moreover, large co-adapted complexes can be inherited from hybridization as opposed to point mutations (Heliconius Sequencing Consortium 2012).

Hybrid populations may have novel trait values outside both of the parent species ranges, and this may allow for adaptation to extreme environments c.f. Rieseberg et al. (1999), or very high trait variability compared to their parents (Lamb and Avise 2016). In general, both additive genetic variation and trait genetic and phenotypic covariances are expected to change when hybridization takes place, altering genetic drift and the response to selection (Bailey et al. 2013; Grant and Grant 1994; Lucek et al. 2016). Lastly, hybridization can have different effects on Linkage Disequilibrium (LD), including increasing or decreasing it, and may hence release genetic constraints (Seehausen et al. 2014). Therefore hybrid species may have different potential to evolve and diversify than their parent species.

Hybrid evolutionary potential (Glossary) can be affected by contingencies in the mosaic of parental genomes arising from the ecological conditions and effective population sizes at initial hybridization (Eroukhmanoff et al. 2013), and the degree of divergence between the parent species at hybridization (Stelkens et al. 2009). Furthermore, evolvability and constraints depend on the mechanisms of divergence among the parent taxa (Rieseberg et al. 1999; Bailey et al. 2013; Chevin and Haller 2014). The involvement of major QTL in species differences (see Glossary) is thought to indicate strong selection during divergence. In contrast, species differences caused by many loci of small effect and the persistence of polymorphism within species are thought to indicate weaker selection, repeated bouts of evolution during divergence, or divergence by genetic drift (Ritchie and Phillips 1998; Orr 2001; Saldamando et al. 2005; Shuker et al. 2005). Major QTL lead to higher trait variation, and hence evolvability, in hybrids compared to their parents due to extra ‘segregational variance’ (Lynch and Walsh 1998). Divergent directional selection among parent species in a multivariate trait (such as plumage colour) would therefore lead to high hybrid segregational variance concentrated along the axis of maximum among-parent differentiation (maximum *Q*_ST_; the discriminant axis), and would facilitate evolution back towards parental phenotypes. However, divergence by stabilizing selection is more likely than directional selection to lead to complementary gene action (Rieseberg et al. 1999; Rieseberg 2003), resulting in higher segregational variance than can be predicted from among-parent differentiation. Thus, stabilizing selection is more likely to result in novel or transgressive trait values in hybrids, and this pattern may not be restricted to axes of high *Q*_ST_ in multivariate traits. Here, we use these expectations to create and examine testable predictions of the impacts of hybridization on the evolutionary process.

We use the Italian sparrow (*Passer italiae*) and its two parent species the house sparrow *P. domesticus* and the Spanish sparrow *P. hispaniolensis* as a study system to address if parental patterns of differentiation are predictive of hybrid species evolution. The Italian sparrow is a homoploid hybrid species distributed on the Italian peninsula and some Mediterranean islands (Summers-Smith 1988; Fig. 1a), mostly in allopatry from its parents. It has an admixed genome intermediate between the parent species, and shows different forms of reproductive isolation with each parent, confirming it as a hybrid species (Hermansen et al. 2011; Elgvin et al. 2011; Trier et al. 2014; Hermansen et al. 2014; Bailey et al. 2015; Elgvin et al. 2017). The house sparrow expanded from the Middle East through the Palearctic region some 3000–7000 years ago, in parallel with the expansion of human agricultural societies (Saetre et al. 2012), and the hybridization event(s) generating the Italian sparrow are thus likely to have occurred thousands of generations ago. The Italian sparrow is sexually dimorphic like its parent species, and male plumage color pattern is intermediate between the two parent taxa (Töpfer 2006; Summers-Smith 1988; Fig. 1b). Moreover, crown color has a narrow geographic cline and is bimodally distributed in a hybrid zone in the Alps between Italian sparrows and house sparrows, suggesting strong selection on this trait (Bailey et al. 2015). We examine crown color and three other sexually dimorphic male plumage color traits in Mediterranean island populations of Italian sparrows, from Corsica, Crete, Sicily and Malta (Fig. 1). These populations are genomically divergent and differ in the proportion of the genome inherited from each parent species, and probably represent independent hybridization events (Runemark et al. 2018).

**Figure 1.**
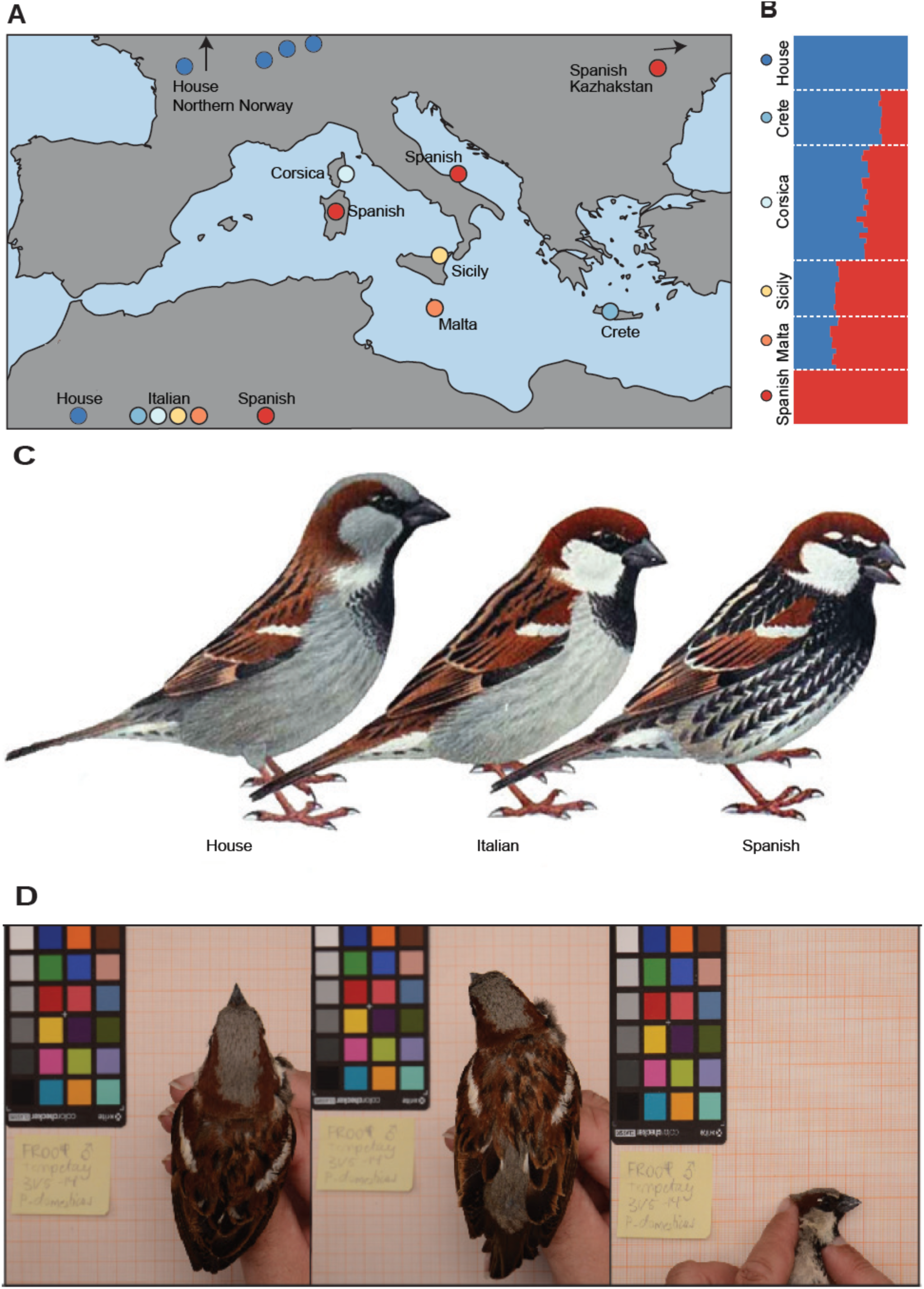
Sampling design, species differences in plumage and study traits. The Italian sparrow is a hybrid between the parent species house sparrow and Spanish sparrow, and originated as the house sparrow expanded eastwards following the spread of agriculture and met and hybridized with island populations of the Spanish sparrow. A) Isolated island populations differing genetically (Runemark et al. 2018) from Crete, Corsica, Sicily and Malta were sampled, as were reference populations of the parent species from France and Switzerland (house sparrows) as well as Sardinia and Kazhakstan (Spanish sparrows). B) The Italian sparrow plumage color pattern is intermediate between the two parent taxa (Töpfer 2006; Summers-Smith 1988). The male has a white cheek and chestnut crown like the Spanish sparrow, but no black flankings. Compared to the house sparrow, the Italian sparrow lacks the gray crown but has the same brown-streaked back, reduced bib and grey rump. C) As males of the Italian sparrow differ from parent species in secondary sexual plumage we chose four plumage traits, crown, back, rump and cheek as study traits. Photographs taken from three different angles were used to extract color information for these characters.

As reproductive isolation from parent species is important for the establishment of homoploid hybrid species (Schumer 2014) and may involve sorting of pre-existing isolating mechanisms in the hybrid (e.g. Hermansen et al. 2014), it is of particular interest to study whether sexually selected traits are constrained to the same combinations in different populations of the hybrid species. Furthermore, diversification of traits involved in mate recognition among hybrid populations may promote further speciation within a hybrid species. We chose male plumage traits as study characters, as plumage is sexually dimorphic in the Italian sparrow and its parent species and hence putatively under sexual selection, and since crown color has been shown to be under strong selection (Bailey et al. 2015). Hence, this variation could be important for premating isolation in the species complex. Using multivariate measures of crown, rump, back, and cheek color (Fig. 1) from the island populations of Italian sparrows and their parent species, we describe patterns of plumage divergence between house and Spanish sparrows, and their relationships with hybrid Italian sparrow phenotypic trait values. We address the current distribution of hybrid phenotypes after many generations of evolution, prevalence of novel phenotypes, divergence among populations and islands, and current evolvability in the Italian sparrow. Based on the expectations outlined in Table 1 we address if parental differentiation can be used to predict hybrid phenotypic evolution.

**Table 1.**
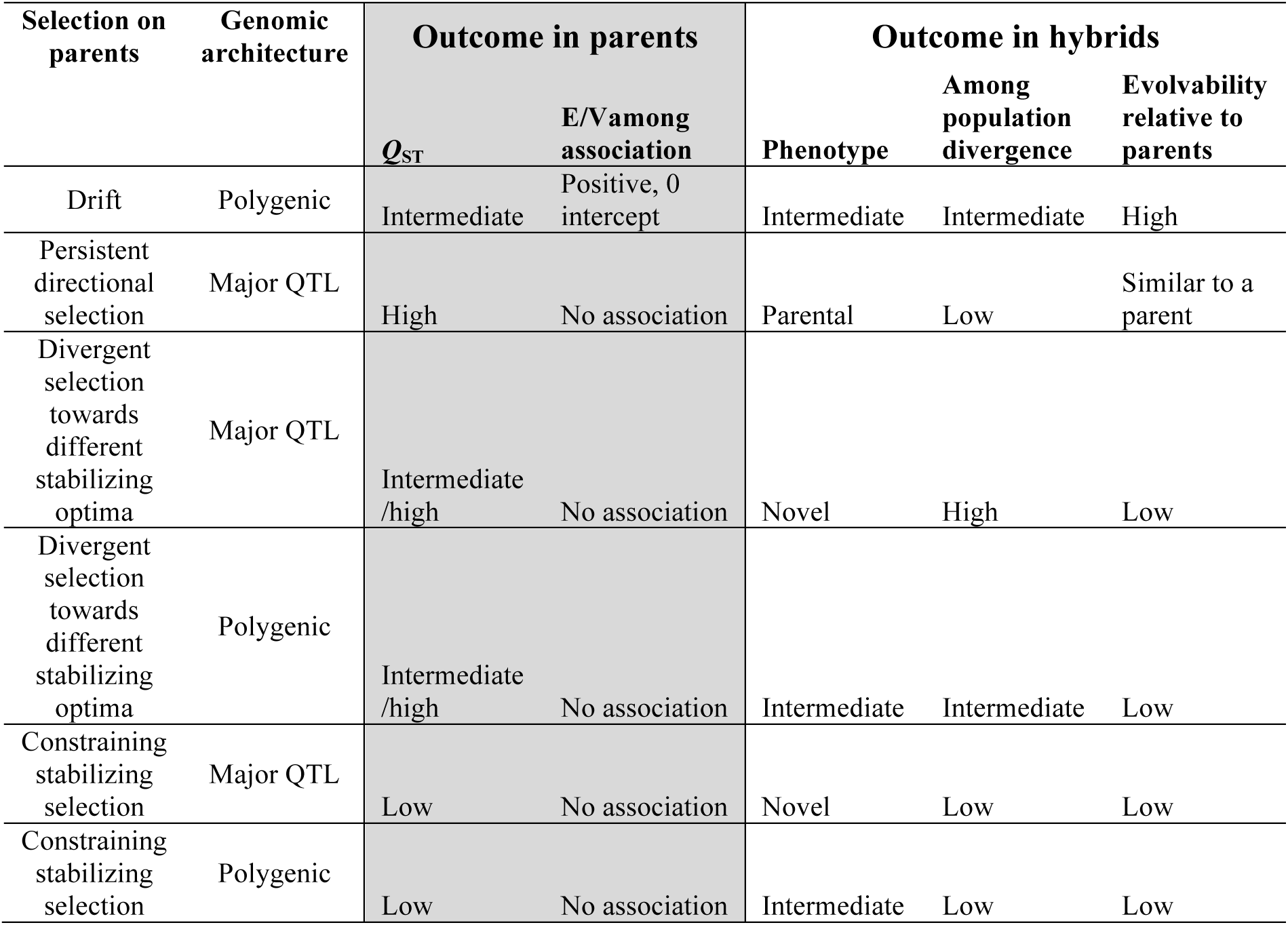
Predictions regarding long-term evolutionary outcome in hybrids given patterns of selection on parent species and genomic architecture. Evolvability relative to parents refers to a hybrid species after many generations of evolution.

| Selection on parents | Genomic architecture | Outcome in parents |  | Outcome in hybrids |  |  |
| --- | --- | --- | --- | --- | --- | --- |
| | | $Q_{ST}$ | E/V among association | Phenotype | Among population divergence | Evolvability relative to parents |
| Drift | Polygenic | Intermediate | Positive, 0 intercept | Intermediate | Intermediate | High |
| Persistent directional selection | Major QTL | High | No association | Parental | Low | Similar to a parent |
| Divergent selection towards different stabilizing optima | Major QTL | Intermediate /high | No association | Novel | High | Low |
| Divergent selection towards different stabilizing optima | Polygenic | Intermediate /high | No association | Intermediate | Intermediate | Low |
| Constraining stabilizing selection | Major QTL | Low | No association | Novel | Low | Low |
| Constraining stabilizing selection | Polygenic | Low | No association | Intermediate | Low | Low |

We compare the four multivariate plumage traits to ask: (1) Have traits differentiated significantly among the three species and the island hybrid populations, and is differentiation increased by island isolation in Italian sparrows? (2) Have traits that are more differentiated between house and Spanish sparrows – suggesting divergent selection – evolved and diverged more among Italian sparrow populations, and have they evolved more strongly back to towards parental trait values? (3) Has each trait evolved more in Italian sparrows along axes of high current parental evolvability, or high parental differentiation, and how are the results related to the above patterns of parental divergence? (4) To what extent do hybrids have novel phenotypes, and can a higher degree of novelty be explained by lower current parental evolvability, suggesting novelty in hybrids caused by fixed differences between parents at major QTL? (5) Combining the four plumage traits, are Italian sparrows less intermediate in phenotype and more a mosaic of parental trait values than expected through evolution by drift? (6) Does present-day hybrid trait evolvability remain higher than the parent species, despite many generations of evolution?

## Materials and Methods

### Sampling

We sampled adult male Italian sparrows from three populations on each of the islands Corsica, Crete, and Sicily during March-June 2013 and on Malta in June 2014. Spanish sparrows were caught on Sardinia in June 2013 and Kazakhstan April 2014 while house sparrows were sampled at two locations in France and one in Switzerland during June 2014 (Fig. 1a; Supporting Table 1). All birds were caught using mist nets, photographed, and released immediately after sampling to minimize stress. We caught, photographed and extracted plumage information for 183 Italian, 53 house and 29 Spanish sparrows. Coordinates for sample sites and the number of photographed individuals per population are summarized in Table S1. All necessary permissions were obtained.

### Plumage color measurement

We photographed the birds in a standardized light environment alongside a color checker (5.7 × 8.7 cm X-rite mini ColorChecker ®classic) using a Nikon D500 (16.2 megapixels) (Fig. S2; for more details about the photographic setup, see Tesaker, 2014). We chose four male plumage traits, crown, cheek, back and rump, for color analysis as these are sexually dimorphic and differ between the parental species (Summers-Smith, 1988; Fig. 1b). We photographed sparrows from dorsal, ventral and lateral angles to cover the plumage areas of interest (Fig. 1c). We used a color quantification method suited to the complex color patterns found in the sparrow (Brydegaard et al. 2012). As each plumage trait itself is multivariate, we used a mean-centered singular value decomposition (equivalent to PCA) as implemented in the Chromatic Spatial Variance Toolbox (Brydegaard et al. 2012) to reduce the between-individual color variation to five to six axes of variation for each trait (Supporting methods, section A). Each of these axes describes an aspect of the within-individual plumage color variation. We hence used four different traits each being multivariate, with 5-6 PC axes (Supporting Table 2, Supporting Figure 2).

**Figure 2.**
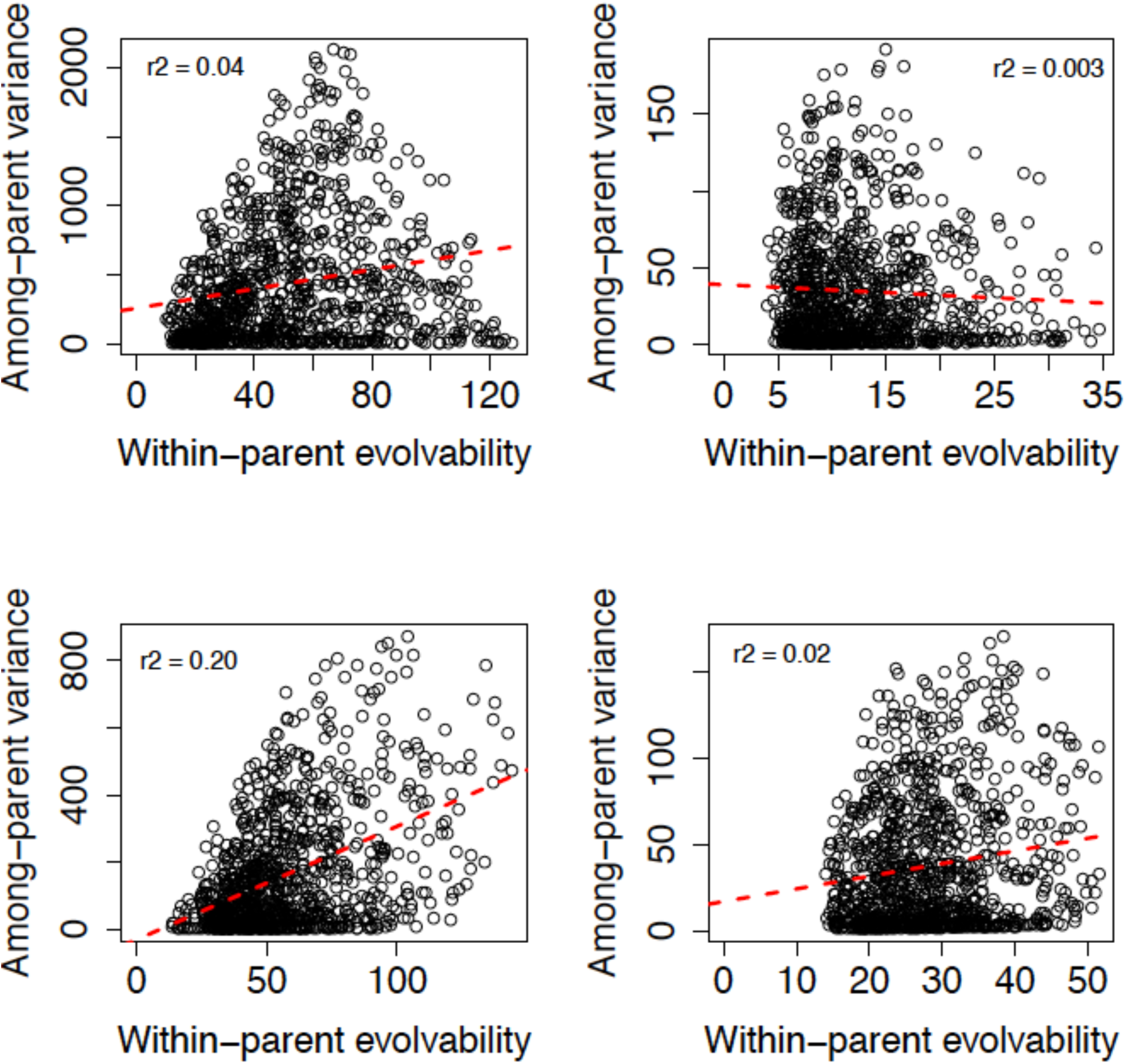
Within- versus among-parent variance along 1000 randomly chosen unit vectors through PC1-5 (crown and cheek) or PC1-6 (back and rump). The red dashed lines are the fitted regression slopes. (a) crown (TOP LEFT), (b) back (TOP RIGHT), (c) rump (BOTTOM LEFT), (d) cheek (BOTTOM RIGHT). The grey dashed lines indicate the mean and variance of a uniform distribution between 0 and 1, and are included for reference only.

### Statistical analysis

To understand the processes shaping hybrid Italian sparrow evolution, we first examined patterns of divergence of our four focal plumage traits in the parent species. In the absence of fixed differences between parent species at major QTL, we expected a positive relationship between current within-parent trait variability and the amount of among-parent trait divergence for each multivariate plumage trait (Schluter 1996; Hansen and Houle 2008; Bolstad et al. 2014).

We then tested whether the magnitude and pattern of parental differentiation was predictive of hybrid Italian sparrow evolution (predictions described in Table 1; Supporting Methods). To this end, we estimated covariance matrices and population trait means for the parent species and the Italian sparrow populations. Specifically, we used PC scores from each of the focal plumage traits to derive (1) a within-population trait covariance matrix for each species; (2) a within-population covariance matrix representing the average of the two parent species (the ‘parental’ covariance matrix, which also represented the predicted F2 generation covariance matrix; see Glossary); (3) trait means for each parent species individually; (4) the average of the parental trait means (representing expected F2 generation trait means, assuming additivity); and (5) the trait means for each individual Italian sparrow population.

In the absence of pedigree information we estimated these based on phenotypic trait values, and hence used phenotypic ‘P’ matrices rather than additive genetic G matrices (see Glossary). P matrices are typically similar in shape to G matrices (Roff 1997), especially for traits with high heritability and low condition-dependence, as can be expected for melanin pigmentation traits (Hill and Brawner 1998). To predict the F2 matrix, we assumed additivity, linkage equilibrium, and among-species variance controlled by many loci of small effect. If there are no major QTL involved in species differences, there is no expected increase in trait variability in a hybrid compared to its parents (i.e. no segregational variance; T. F. Hansen pers. comm.; see Lynch and Walsh 1998 page 228, equation 9.17, for the univariate expectation). We therefore estimated the predicted F2 matrix as the average of the parental P matrices. F2 mean trait values were also predicted to be the average of the parents, as we assumed additivity.

To estimate the covariance matrices and population means we created posterior distributions from a model with population as fixed effect using the Bayesian mixed model R package MCMCglmm (Hadfield 2010). This approach allows uncertainty to be taken into account in downstream analyses (Hadfield 2010). For Italian sparrow within-population values we removed the intercept (‘fixed = trait:Pop – 1’ giving trait mean estimates per population) and estimated the full residual covariance matrix (‘rcov = ∼ us(trait):units’). The same model structure was used to estimate the within-population (residual) covariance matrix for each parent species. To estimate each parent species’ trait means, the model was the same except that ‘population’ was instead included as a random effect (‘fixed = trait – 1’ giving the trait means; ‘random = ∼ idh(trait):Pop’). For all models, default priors were used for fixed effects, and a diagonal matrix of variance 1 was used as a prior for residuals and random effects where appropriate (e.g. ‘V = diag(6), n = 7’ for plumage traits with 6 trait variables; ‘V = diag(5), n = 6’ for 5 variables). All plumage trait values (PC axis scores for 5 or 6 axes per trait) were multiplied by 100 to increase trait variances above 1. This does not affect our conclusions, which are not based on absolute variance values. All models were run for 100k iterations including 10k burnin, and post-burnin thinning of 50, making 1800 posterior values per parameter.

#### Among-parent divergence

We tested whether parental trait divergence was explained by current within-parent evolvability, using the average of the parental P matrices for the latter. Using the parental means and covariance matrix, we estimated within-species evolvability (eB, averaged for the two parent species; Hansen & Houle 2008; Bolstad et al. 2014), among-species variance (V_among_), and among-species *Q*_ST_ as V_among_/(V_among_ + 2eB) along each of 1000 randomly selected axes (unit vectors) through multidimensional trait space (created in the R package ‘evolvability’; Bolstad et al. 2014) plus the linear discriminant axis, which represents the axis of maximum *Q*_ST_ (see Glossary). This approach uses the hypothesis that fixed major QTL for species differences would reduce this association by reducing current evolvability relative to divergence along some axes, and such QTL would indicate strong selection during divergence. In the absence of strong selection, we expect among-population variance to scale to within-population evolvability along each axis of variation (analogous to Martin et al. 2008). First we regressed among-parent variance (response) on average within-parent evolvability (predictor) along the 1000 randomly chosen unit vectors (Bolstad et al. 2014; Schluter 1996) for each trait. In the absence of major QTL we expected a positive slope, a high proportion of variance explained (*R*^2^), and an intercept close to zero. We then compared the average, variance and maximum among-parent *Q*_ST_ among traits along the same 1000 random axes plus the parental discriminant axis (which has highest *Q*_ST_; see Glossary) to assess evidence for differences in the strength of divergent selection among traits, and the potential influence of stabilizing selection on parental divergence (Table 1).

#### Italian sparrow phenotypic trait values, among population divergence and evolvability

We assessed relationships between the magnitude and pattern of parental divergence described above, and evolution in the Italian sparrow. In the absence of major QTL, we expect hybrid evolvability to be similar to current parental evolvability, but in their presence we expect increased variation in the hybrid along axes of high parental differentiation and/or low parental evolvability.

#### Hybrid trait values

We tested if hybrid phenotypes tended to be distributed along axes of high within-parent evolvability or high among parent *Q*_ST_. To do this, we first estimated the axis of divergence of each individual’s trait value from the expected F2 means by calculating the loadings of the plumage PC scores (Supporting Table 2) along that axis using the method of Schluter (1996). We tested whether each of among-parent *Q*_ST_ and average within-parent evolvability differed significantly between this set of individual axes compared to 1000 random axes, using non-overlap of the 95% Highest Posterior Densities (HPD) of the two data sets as a significance test. If phenotypes tended to be distributed along axes of high parental evolvability in traits with moderate or low parental divergence and strong relationships between within- and among-parent variability (above), this would support neutral or weakly selected hybrid and parental evolution. If hybrid phenotypes were biased towards axes of high among-parent *Q*_ST_ for traits with high parental divergence and weaker variability relationships, this would suggest that stronger selection and major QTL were important for these traits.

#### Hybrid potential for novelty

To investigate hybrid novelty, i.e. if hybrid individual phenotypic traits are outside the ranges of either parent, we first calculated the score of each individual along its axis of divergence from predicted F2 mean trait values (Supporting Figure 3; Glossary), using the loadings as above. We then calculated the score along the same set of axes of the upper and lower 2.5% quantiles for each parent species’ trait values. The novelty index (see Glossary) was the lowest value of ‘individual score – parental quantile score’ among the four parental quantiles. Zero was considered the threshold above which the individual’s phenotype lies outside those expected for either parent species.

**Figure 3.**
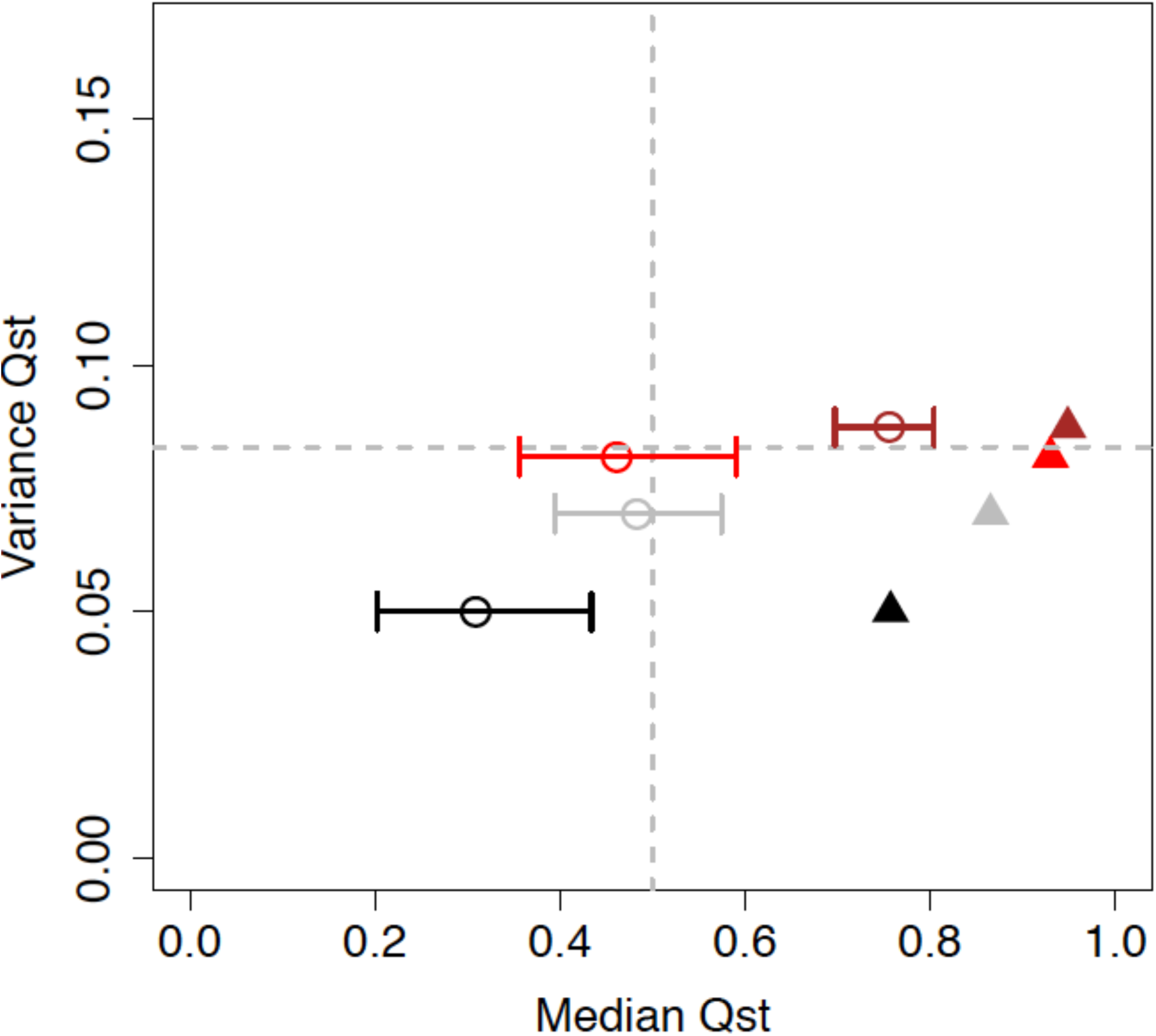
Median, variance and maximum among-parent Qst along 1000 randomly chosen unit vectors for each trait: crown (brown), back (red), rump (grey, expected to most closely match a trait diverging by drift), and cheek (black). Error bars show the upper and lower Highest Posterior Density. Filled triangles indicate maximum Qst values (i.e. Qst of the discriminant axis) on the X axis.

To test if novelty was higher along axes of lower current parental evolvability, indicating major QTL in species differences, we carried out multiple linear regressions on novelty index for each individual along its own axis of divergence from the expected F2 values (see Glossary; response variable), against both parental evolvability and among-parent *Q*_ST_ along the same axes. *Q*_ST_ was included because novelty is expected to decrease with increasing *Q*_ST_ for all traits, as parental means get further apart relative to evolvability, and this would confound relationships between hybrid novelty and parental evolvability. For each trait we also recorded the maximum novelty index and the proportion of novel hybrid phenotypes, as these were expected to differ depending on parental divergence mechanisms (Table 1).

#### Hybrid among-population divergence

For each plumage trait, we first performed a MANOVA to test if species and island populations of the Italian sparrow were significantly different using all PC axes (Supporting Table 2), followed by ANOVA on individual plumage PC axes and univariate post hoc TukeyHSD tests for differences between groups. We also carried out Canonical Variates Analysis (CVA) for multivariate post hoc comparisons of islands and species, applying Bonferroni corrections to P values from 10 000 randomly permuted data sets. For the CVA we used the ‘morpho’ package in R (Schlager 2013).

#### Impact of island isolation on among-population divergence

To test for an effect of among-island isolation on population divergence in a hybrid species, we calculated the average ratio of among-island *Q*_ST_ (*Q*_ST_ of island means) to among-population *Q*_ST_ (*Q*_ST_ of pooled, island mean-centered, population means) across 1000 random axes. If the median ratio is higher than 1 and HPD intervals do not overlap with 1, it means that island isolation promotes divergence.

#### Evolution along the parental discriminant axis

We tested whether divergence among Italian sparrow populations was greater than average along the discriminant axis, as expected if both parents are close to their optima and hence selection drives hybrid populations towards both parental phenotypes (Supporting Methods). We did so by calculating and comparing the median and 95% HPD of variance of population F2 quantiles (see Glossary) on average across 1000 random axes with among-population variance in F2 quantiles along the parental discriminant axis (Glossary). To identify evolution towards either of the parent species, we calculated population F2 quantiles along the parental discriminant axis, with values >0.5 indicating evolution towards Spanish sparrows, whereas <0.5 indicated evolution towards house sparrows.

#### Mosaicism

By combining all four plumage traits, we looked for evidence for higher phenotypic mosaicism (hybrids having a mosaic of parental trait values for the four traits, rather than each trait being intermediate in phenotype) than expected by chance. The rationale for this test is that selection back towards different parental phenotypes in different plumage traits would produce a more mosaic phenotype than neutral expectations. The test was performed first by calculating the mean among-parent discriminant axis score for each Italian sparrow population for all four plumage traits combined, i.e. how similar the Italian sparrow population overall is to the parent species on average. We compared these means to a neutral expectation based on genetic drift (see below). To do this, we first calculated the discriminant axis score of the predicted F2 trait means, and the predicted F2 variance along that axis, for each trait. We then estimated the F2 quantile for each Italian sparrow population mean score, reflecting the distance from the predicted F2 values along the discriminant axis taking into account the estimated variation of the F2 values. Such quantiles were calculated for each trait. The mean and variance of these quantiles across the four plumage traits (crown, back, rump, cheek) were then calculated for each population. We used the variance as the measure of population mosaicism, and compared these to neutral expectations for a given mean. For neutral expectations, we simulated 10k sets of 4 values from a uniform distribution between 0 and 1 (representing random values for the four trait F2 quantiles) and calculated the means of all 10k samples, followed by the variance within each 0.1 unit subset of the mean values. We then tested if there were significant differences between actual trait mosaicism values and neutral expectations for each population.

#### Hybrid population evolvability

The ability of a population to respond to selection, averaged across directions in trait space, can be estimated as its average evolvability (Hansen and Houle 2008). Using the species-level trait covariance matrices, each species’ average evolvability and 95% Highest Posterior Density (HPD) was estimated as the average of evolvability along the 1000 random axes, using the ‘evolvabilityBetaMCMC’ function in the evolvability package (Bolstad et al. 2014). Evolvability is expected to drop slowly and still potentially be high after many generations in a hybrid species evolving under drift, be lower under strong stabilizing selection, or be similar to one of the parent species if there is persistent directional selection towards that phenotype (Table 1).

All analyses were performed in R v. 3.3.1 statistical software (https://www.r-project.org/). For the majority of analyses (except ANOVA, MANOVA and CVA) we used a modified version of the ‘evolvabilityBetaMCMC’ function from the R package ‘evolvability’ (Bolstad et al. 2014) to calculate quantities based on the full MCMCglmm posterior distribution of trait means and covariance matrices. For further details, see the annotated code in Github (XXXX).

## Results

### Patterns of among-parent divergence and inferred selection on parental traits

In the absence of fixed major QTL causing species differences, evolvability within and between groups is expected to be correlated. The rump was the trait for which within- versus among-parent variance relationships matched most closely (Table 1; Supporting Table 3). The proportion of among-parent variance explained by within-parent evolvability in rump color was at least an order of magnitude higher than for the other traits (Figure 2; Supporting Table 3; rump *R*^2^=0.20). All slopes except for back color were highly significantly positive, and all intercepts were highly significantly different from zero except rump, which was marginally significant (Supporting Table 3).

Based on this, we expect the average, variance and maximum of among-parent *Q*_ST_ for rump color to best represent those of a trait evolving in the absence of strong selection, and assessed differences between rump and other traits to infer what selection pressures are likely to have been acting on the other traits. Crown color had significantly higher average *Q*_ST_ than rump and all other traits, the highest maximum *Q*_ST_ of all traits, and the highest variance, and was therefore the trait most likely to have evolved under divergent selection (Figure 3). Back color had a similarly high maximum suggesting divergent selection along some axes, but the average *Q*_ST_ was much lower, implying that some axes were constrained to low divergence, possibly by stabilizing selection. Back color *Q*_ST_ variance was higher than that for rump, hence increasing the likelihood that a combination of forces had been acting on this trait. Cheek color had the lowest average, maximum and variance in *Q*_ST_, suggesting that divergence may be constrained in this trait. We used these inferred patterns of selection in the parents to test if they are predictive of the patterns for hybrid evolution, using the expectations outlined in Table 1.

### Hybrid trait values

Traits not subject to strong selection in parent species are expected to evolve along axes of largest current parental variation in the hybrids, whereas traits under divergent selection are expected to evolve along axes of high among-parent *Q*_ST_, and traits or axes under stabilizing selection are expected to have more novel hybrid trait values (Table 1). As expected for a trait potentially diverging by drift or weak selection in the parent species, rump was the only trait with hybrid phenotypes distributed along axes with significantly higher average within-parent evolvability than expected by chance (Supporting Table 4). Crown and back color were inferred to be partly under divergent selection in parent species and did, as predicted, have hybrid trait values that were significantly biased towards axes of high among-parent *Q*_ST_ (Supporting Table 4).

### Hybrid potential for novelty

The highest novelty is expected for traits in which parental trait values have been under stabilizing selection, causing low phenotypic relative to genetic divergence. Interestingly, the amount of novelty along each individual’s axis of divergence from the predicted F2 means was high for all traits, with the highest proportion for back color, where stabilizing selection on some color dimensions in the parent species are inferred (0.64), followed by rump (0.55), crown (0.54) and cheek (0.42); see Supporting Figure 4. These patterns differed with island of origin. While Crete generally had high novelty, Corsica had low, Sicily and Malta both had high novelty for back, Sicily had novel rumps, and Malta had novel crowns (Supporting Figure 5; Supporting Table 5).

**Figure 4.**
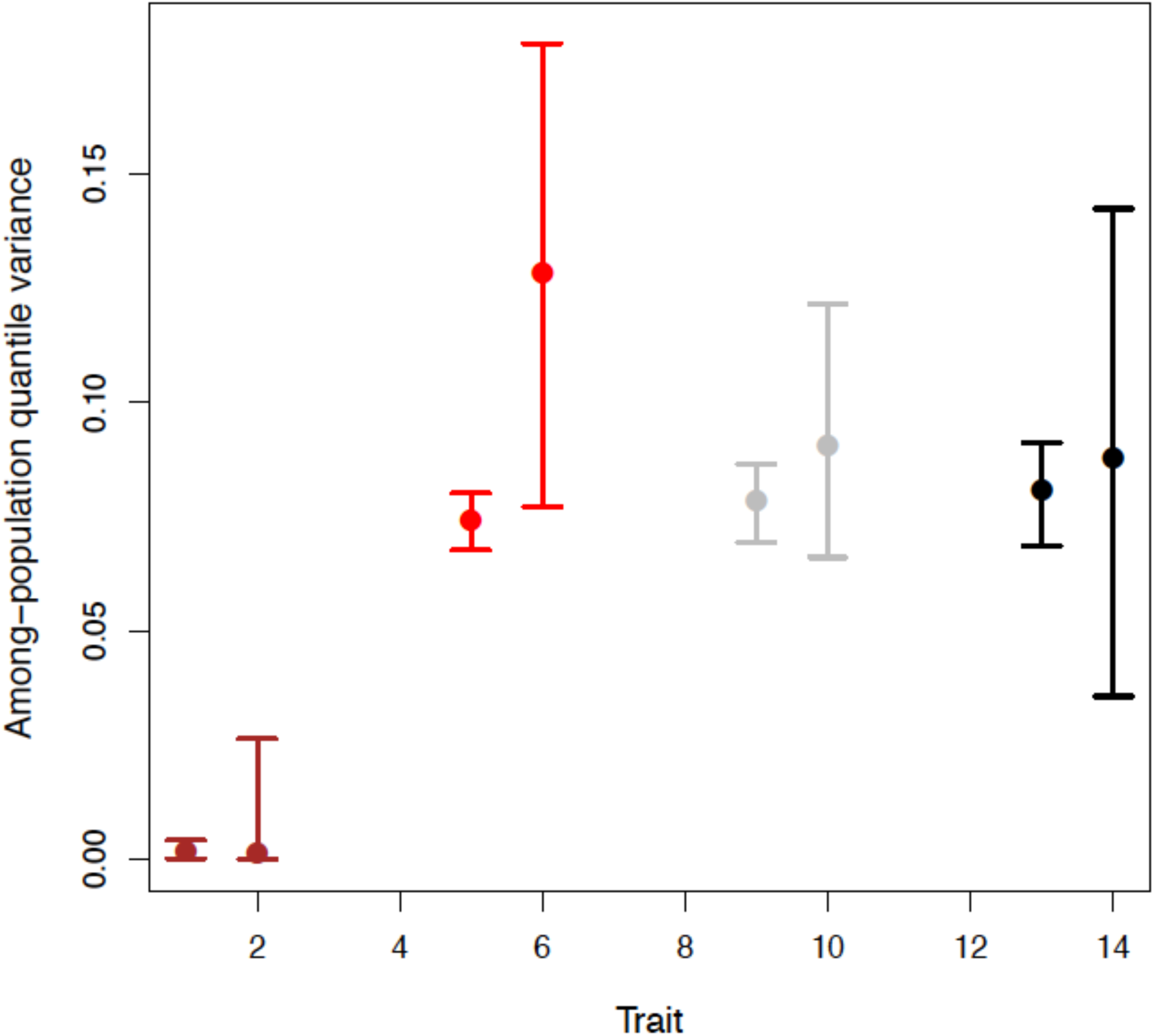
Among-population (not distinguishing within- versus among-island) variance in F2 quantiles of population means for crown (brown bars), back (red), rump (grey) and cheek (black). The left bar is the median and 95% HPD of 1000 random axes (which are mostly partially correlated with the discriminant axis, reducing the likelihood of a significant difference), and the right bar is the median and HPD of the discriminant axis among-population variance.

**Figure 5.**
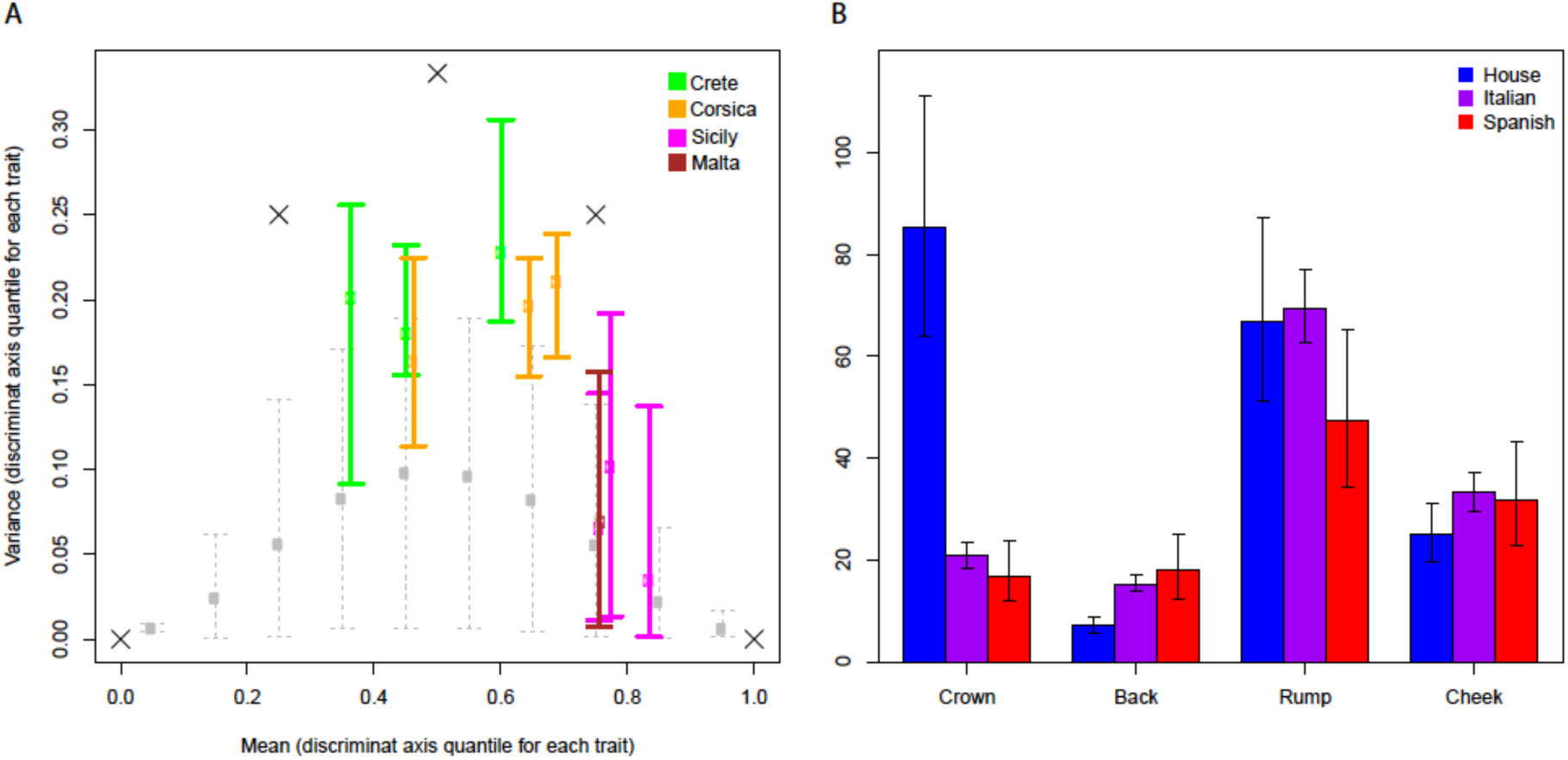
Plumage mosaicism and evolvability. A) Plumage mosaicism in Italian sparrows. Each colored point is an Italian sparrow population (Table S1; green = Crete, orange = Corsica, magenta = Sicily, brown = Malta). The x axis is the mean (across the four plumage traits) of the F2 quantile of the population trait mean. The y axis is the variance of the same quantity and represents the measure of mosaicism. Error bars for population variance are the 95% highest posterior density (HPD) given the MCMCglmm posterior distribution. The grey points and dashed error bars represent the mean and 95% HPD of neutral expected variance within each 0.1 unit subset of the mean values, for the 10k simulated neutral variance values. The large black crosses represent the maximum possible mosaicism for mean = 0, 0.25, 0.5, 0.75 and 1 (cases where all four trait quantiles are either 0 or 1). B) Evolvability in the Italian sparrow and its parent species. Bars denote means, and error bars denote 95% HPD, parent species are illustrated in blue (house sparrow) and red (Spanish sparrow) and the Italian sparrow illustrated in purple.

Controlling for confounding variation in among-parent *Q*_ST_, the relationship between parental evolvability and hybrid novelty was significantly negative for all traits except for rump (Supporting Table 6), indicating that for crown, cheek and back, low parental evolvability may be due to fixation of major QTL.

### Divergence among hybrid populations

The parent species and all individual islands were significantly different from each other for all dimensions and all traits combined, with Corsica and Crete being the most similar (Supporting Tables 7-8). For crown color, the grey crowns of the house sparrows were significantly different from the brown crowns of Italian and Spanish sparrows. The only other significant difference for crown color was between Corsica and Crete indicating that, overall, Italian sparrows are consistently very similar to Spanish sparrows for this trait. Corsica and Crete did not differ significantly from house sparrows in back plumage, but all other comparisons were significant, indicating strong divergence among species and islands for this trait. Rump color showed similarly high levels of diversification, with the only non-significant comparisons being between Corsica and Crete, and Malta and Spanish sparrows. Cheek color was less divergent than back and rump, with complex relationships among islands and species. In species-level comparisons, Italian sparrow crown and cheek color were not significantly different from Spanish sparrows. All MANOVA analyses of all axes for individual traits revealed highly significant differences between species and islands (Supporting Table 9). For all traits except cheek, Italian sparrows were significantly different from each parent species along at least one PC axis (for cheek they were never significantly different from Spanish sparrows; Supporting Table 10).

### Impact of island isolation on among-population divergence

For all traits except cheek, island isolation had a significant impact on divergence, as the average ratio across 1 000 random axes of among-island to among-population (within island) *Q*_ST_ was significantly higher than 1 (Supporting Table 11). This also confirms that islands have evolved in different directions (Supporting Table 11). Back, rump and cheek had remarkably similar average among-population *Q*_ST_, while crown had significantly lower among-population *Q*_ST_, indicating that low heritable variation is preventing the buildup of differentiation by drift (Supporting Table 11). Therefore hybridization can lead to multiple different phenotypes, especially among isolated populations.

### Evolution along the parental discriminant axis

The low hybrid *Q*_ST_ of crown color supported uniform and persistent directional selection having caused reduced polymorphism in this trait. This was further supported by crown being the only trait with extreme median discriminant axis F2 quantile across all populations, with all populations having evolved strongly towards Spanish sparrows (Supporting Table 12). The best evidence for the action of divergent selection in the hybrid was for back plumage. Crown and cheek had lowest among-island *Q*_ST_, with rump *Q*_ST_ significantly higher than both, and back *Q*_ST_ significantly higher than rump (Supporting Table 12). Back also had highest average island:population *Q*_ST_ ratio (i.e. relatively more among-than within-island differentiation than the other traits), significantly higher than cheek and rump (see previous paragraph).

Furthermore, back was the only trait with marginally higher *Q*_ST_ (not differentiating island- and population-level) along the discriminant axis than 1 000 random axes (Figure 4), indicating that much of the divergence was along the primary axis of parental differentiation. This is consistent with evolution back towards fit parental phenotypes for axes most likely to have undergone divergent selection among parents.

### Mosaicism

If selection would favour the hybrid species to diverge towards the phenotype of one of the parent species, lower mosaicism than the random expectation would be predicted. However, none of the population mean mosaicism values were below the nearest neutral mean values (Figure 5a). Mosaicism values above neutral expectation suggest selection towards different parent species in different traits. Corsica and Crete showed the best evidence of an influence of selection causing among-trait mosaicism of parental phenotypes, with the 95% HPD of two populations falling fully above those of the neutral expectations, and none of the population HPD intervals overlapping with the mean neutral expectation (Figure 5a).

### Hybrid population evolvability

Italian sparrow evolvability was not significantly higher than that of the parents for any of the plumage traits, although the mean was slightly higher for rump and cheek (Fig. 5b). As the Italian sparrow crowns have evolved back towards a Spanish phenotype, we predict Italian sparrow crown plumage to have similar evolvability to Spanish, and it does (Fig. 5b). The high evolvability for house sparrow crown is probably caused by a number of the house sparrows that had some brown in the crown.

## Discussion

While there is evidence that hybridization can alter the covariance between traits (Lucek et al. 2016), the effect of hybridization on individual traits is not well understood. Although there are theoretical expectations based on the dispersion of mutations predicted under different types of selection on the parent species (Bailey et al. 2013), these have not been empirically tested. Here, we show that patterns of divergence and inferred selection on parental phenotype are indeed predictive of trait variation in the hybrid Italian sparrow, and move the research field from a description of patterns of divergence closer to addressing the processes involved.

Our findings match predictions based on parental patterns of divergence surprisingly well, supporting the validity of our approach for inferring selection in the parent species based on an expected presence of major QTL for species differences when selection is strong. As predicted, traits putatively subject to weak selection or drift in parental house and Spanish sparrows evolve along axes of high current within-parent evolvability in hybrid Italian sparrows. Traits putatively under stronger selection are biased towards axes of high among parent *Q*_ST_, and those under stabilizing selection show transgression (see Glossary) in hybrids. Interestingly, we also find a negative relationship between parental evolvability and hybrid novelty, suggesting that there may be fixed major QTL differentiating parent species at the genomic level whether or not there is strong phenotypic divergence. Axes with low parental evolvability are expected to be maintained at low mutational variance by strong drift or stabilizing selection, and be most affected by complementary gene action acting on major QTL.

Using traits under different selective regimes in parent species, as in the present study, increases our ability to test predictions. However, while we have inferred selection on the parent species, we only have previous independent evidence for selection on crown color (Bailey et al. 2015). Although sexual dimorphism in all study traits implies that the traits are likely to be under some kind of sexual selection, caution is needed when interpreting the results. As a high proportion of homoploid hybrid animal species live in sympatry with at least one parent species (Mavárez and Linares 2008), differentiation in sexually selected traits is important for maintaining the integrity of hybrid species through assortative mating (Schwander et al. 2008; Melo et al. 2009). The potential for such assortative mating depends on trait values of sexually selected traits and the preferences for these. Interestingly, secondary sexual traits may be under different forms of selection. ‘Species recognition’ traits are expected to be under stabilizing selection, whereas adaptive mate choice traits indicating vigor may be under consistent directional selection (Bentsen et al. 2006). Novel trait values also increase the likelihood for the establishment of pre-mating isolation (Doherty and Gerhardt 1983; Gompert et al. 2006; Magalhaes and Seehausen 2010). If our findings of higher potential for novelty in traits putatively under stabilizing selection (e.g. back color) are general, this would increase the likelihood of transgressive values for species recognition traits and hence facilitate the establishment of pre-zygotic isolation.

Predicting how hybridization will influence evolution and the response to selection will increase our understanding of its role in evolution. For instance, animal hybrid species are typically ecologically divergent from both parental species. They may occupy different habitats requiring new adaptations (Nolte et al. 2005; Gompert et al. 2006; Nice et al. 2013). Transgression could enable the use of a niche that is distinct from parental species’, which theoretical models predict to be important for successful establishment of a hybrid species (Buerkle 2000; Duenez-Guzman et al. 2009). Hence, not only the ecological context in which hybridization occurs, which has been suggested to be important (Buerkle 2000; Mallet 2007), but also patterns of selection on parent species for key ecological traits may be crucial for successful establishment of hybrid species. The potential for this is likely to be highest when parent species are under stabilizing selection for key ecological traits since theoretical predictions (Bailey et al. 2013; Rieseberg et al. 1999) and our results suggest they have a high potential for transgression in hybrids, e.g. back color in this study.

Transgression in individual characters is not required for pre-zygotic isolation to establish. If sexually selected traits are determined by several genes, hybridization may recombine features of both parents into new trait combinations (Doherty and Gerhardt 1983; Mavárez et al. 2006; Carlos et al. 2009; Hermansen et al. 2014) that may differentiate hybrid species from both their parents. We find evidence for such mosaic inheritance of traits, providing a novel differentiated phenotype, within the Italian sparrow. This unique trait combination may facilitate the co-existence with the Spanish sparrow in the Gargano peninsula without introgression (Sætre et al. 2017). Trait mosaicism is strongest in the Crete and Corsican populations, where variance among traits in their value along the discriminant axis between parent species is significantly higher than expected by chance. Interestingly, the Sicilian and Maltese birds are closer to the Spanish sparrow along the discriminant axis, and variance for these populations is not higher than expected by chance. Potentially, these populations could have undergone unidirectional backcrossing, e.g. due to purging of incompatibilities, c.f. (Baack and Rieseberg 2007). Whole genome data is consistent with this scenario, as the proportion of the genome inherited from the Spanish sparrow is approximately 0.75 for the Sicilian and Maltese populations (Runemark et al. 2018).

Elevated evolvability due to the high polymorphism may be expected in early generation hybrids, but we find no evidence for significantly increased evolvability in current Italian sparrow populations. As the hybridization event(s) leading to the Italian sparrow are thought to have taken place thousands of years ago (Hermansen et al. 2011; Saetre et al. 2012) the study populations may evolved and lost variation by drift and selection. Whether increased evolvability is maintained may depend on whether drift reduces polymorphism down to the same level as at mutation-drift balance in the parents. For traits that have evolved strongly back towards one parent, evolvability is expected to be similar, which is consistent with our findings for crown.

The significant among-island diversification provides another interesting perspective of the potential of hybridization to create diversity. Potentially, the diversification from different mosaic combinations of parental traits we find may mean that reproductive isolation can arise between lineages originating from hybridization between the same parental species.

Sexually selected traits, and the preferences for them, may co-evolve along a line of equilibrium where the trait matches the preference in isolated populations (Uyeda et al. 2009), and this may be part of the explanation to the diverse combinations of plumage traits in the isolated island populations in this study. Ecologically selected traits are expected to evolve in response to local environment e.g. (Runemark et al. 2015). Interestingly, we find evidence for beak size being adapted to annual temperature in the same island populations as in this study (Runemark et al. *in press*). While beak size seems to easily evolve and is best explained by local temperature and precipitation patterns, consistent with patterns from the fossil record (Hunt 2007), we find some evidence consistent with genomic contingencies in form of a significant effect of genomic similarity to a parent species on similarity to that parent species in beak shape (Runemark et al. *in press*). Such contingencies may also be important for plumage divergence.

### Methodological considerations

In this study, a phenotypic P matrix was used to study the evolutionary potential of a hybrid species, when commonly researchers use an additive genetic variance matrix, or G matrix. Although the phenotypic correlations might not accurately reflect the genetic correlations, increasing evidence shows that the differences between phenotypic correlations and genetic correlations are minor enough for a P matrix to roughly represent a G matrix (tested in Cheverud 1988; Roff 1995; Steppan 1997; Waitt and Levin 1998). Further, heritability of plumage traits may explain the type of selection acting on that particular trait, and we have no direct heritability estimates for these traits. Melanin-based plumage traits, as in sparrows, are predicted to be highly heritable (Hill and Brawner 1998), while carotenoid based colors are more condition-dependent (Hill and Montgomery 1994). Sparrows do not have any carotenoid-based colors, which reduces the likelihood that the traits analyzed here are condition-dependent.

The method for used for inferring the strength of selection acting on parent populations is new and previously untested. While phylogenetic tests that examine phenotypic trait variation within and among species along an independently derived phylogeny can be used to distinguish selection pressures (e.g. Eng et al. 2009; Thomas and Freckleton 2011; Kutsukake and Innan 2012), these are not applicable for only two parent species. An alternative is *Q*_ST_-*F*_ST_ comparisons, but these require *F*_ST_ data. Instead, we use a novel approach where we expect that the among-species variance scales to the within-population covariance matrix, in the absence of strong selection causing major QTL in species differences. This is analogous to the expectation that the means of a set of populations diverging by drift should be distributed according to their ancestral population’s G matrix (Martin et al. 2008). Some assumptions need to be made to infer selection on parental species with this approach. With respect to divergence between parental house and Spanish sparrows, the prediction is complicated by the fact that it is based on the G matrix of an unknown ancestral population. We used the average of the parental P matrices as a surrogate for this ancestral matrix. Moreover, the expected pattern of Martin et al. is only the average expectation among many drifting populations, and may not apply to particular pairwise combinations, and finally, the G matrix itself should evolve by drift (Phillips et al. 2001). On the other hand, while Phillips et al. noted large variation in the G matrices of many bottlenecked *Drosophila* populations from the same source, the majority still clustered around the average. Furthermore, as yet we have no specific knowledge of whether the ancestors of house and Spanish sparrows were subject to severe bottlenecks, without which the G matrix may be more consistent across populations. Despite potential limitations and a lack of published information on the expected error structure of population and G matrix divergence, we were nevertheless able to predict hybrid evolution based on parental evolution. If all traits in the sparrows were in fact evolving by drift, or selection did not lead to our expected pairwise relationships, deviations from expectations would be too unpredictable and we would find little relationship between parental divergence and hybrid evolution and evolvability.

### Conclusions and significance

Our findings of predictable outcomes from comparisons between parental divergence and hybrid evolution are directly relevant to the understanding of how hybridization can create novel variation for selection to work on. Empirical demonstrations of successful predictions of patterns also imply that the processes generating the variation are better understood. Predicting outcomes of hybridization is increasingly needed for management, as closely related species come into contact following global warming and range expansions.

## Glossary

Explanations of terms and technical expressions. Terms applicable in the current study only are indicated by an asterisk.

Term: **Explanation**
QST: *Q*_ST_ is a measure of the amount of genetic variance among populations relative to the total genetic variance in the trait (quantitative genetic analogue of FST)
Evolvability: A measure the ability of a population to evolve in the direction of selection when stabilizing selection in other directions is absent. Measured as the response in one generation to unit selection, and equivalent to the additive genetic variance along that axis.
Transgressive segregation (transgression): Extreme trait variance leading to the production of phenotypes that lie outside the parental range
Evolutionary potential: The ability of a population to respond to natural or artificial selection pressures
Genotype variance-covariance (G) matrix: A G-matrix is a summary of the genetic relationships among a suite of traits – their additive genetic covariance matrix – and is central to understanding the ability of multivariate traits to respond to selection
Phenotype variance-covariance (P) matrix: A P-matrix is the phenotypic equivalent to the G-matrix and summarizes the phenotypic relationships among a suite of traits
Quantitative Trait Locus (QTL): A QTL is a locus or genomic region that correlates with variation in a phenotype, potentially harbouring the genes coding for the trait
*F2 hybrid P matrix: The predicted P matrix for an F2 generated from the two focal populations or species. Assuming linkage equilibrium and many loci of small effect governing species differences, this is expected to be the average of the parental P matrices.
*Parental discriminant axis: The linear discriminant axis – which is the axis of highest Q_ST_ and hence greatest differentiation – between parent species in multivariate space.
*Axis of divergence from the expected F2 means: The direction in multivariate trait space of divergence between the predicted F2 means (average of parental means) and an individual or population.
*Novelty index: The difference between the magnitude of divergence of an individual or population from the predicted F2 means, and the magnitude of difference between F2 means and the furthest 97.5% quantile of parental values, along the same axis. Positive values indicate a novel phenotype, see Supporting Figure 3.
*Population F2 quantiles: The quantile (position on the normal distribution, from 0 to 1) of a population trait mean with respect to the predicted F2 mean and variance along either its axis of divergence from the F2 or the among-parent discriminant axis. It is used as a measure of how far a population has evolved from the predicted F2 mean relative to predicted F2 evolvability.
*F2 quantiles along the discriminant axis: As for Population F2 quantiles, but measured along the among-parent discriminant axis rather than the axis of divergence of the population from the F2. Values below 0.5 indicate evolution towards house sparrow phenotypes; above 0.5 towards Spanish sparrows.

## SUPPORTING INFORMATION

**Supplementary Figure 1.**
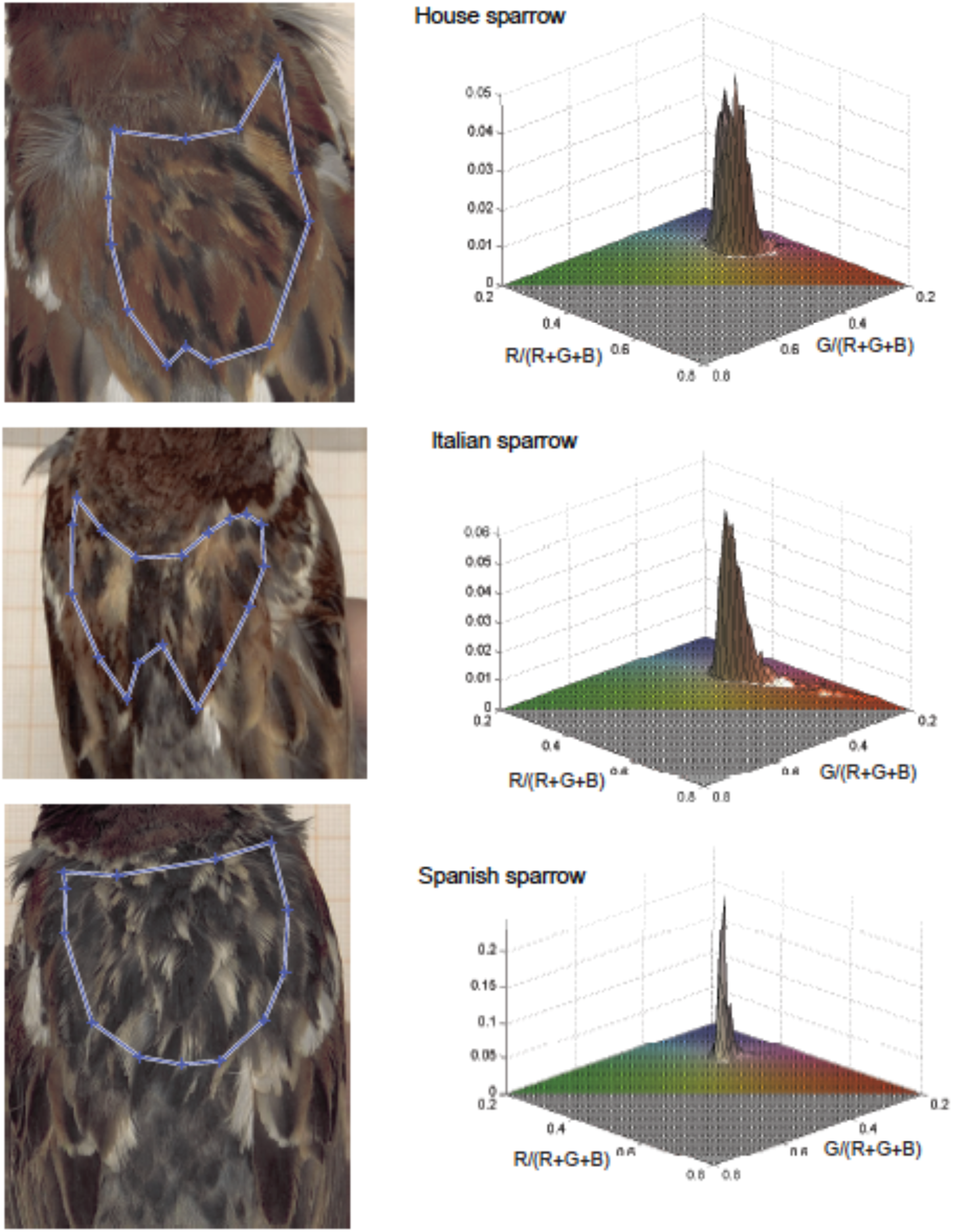
Illustration of the 2D histograms of pixels positioned in RGB-space, capturing the full variance in coloration. These are the raw data the Singular Value Decomposition color analyses for crown, back and rump build on, whereas 3D histograms, incorporating reflectance, are used for cheek.

**Supplementary Figure 2.**
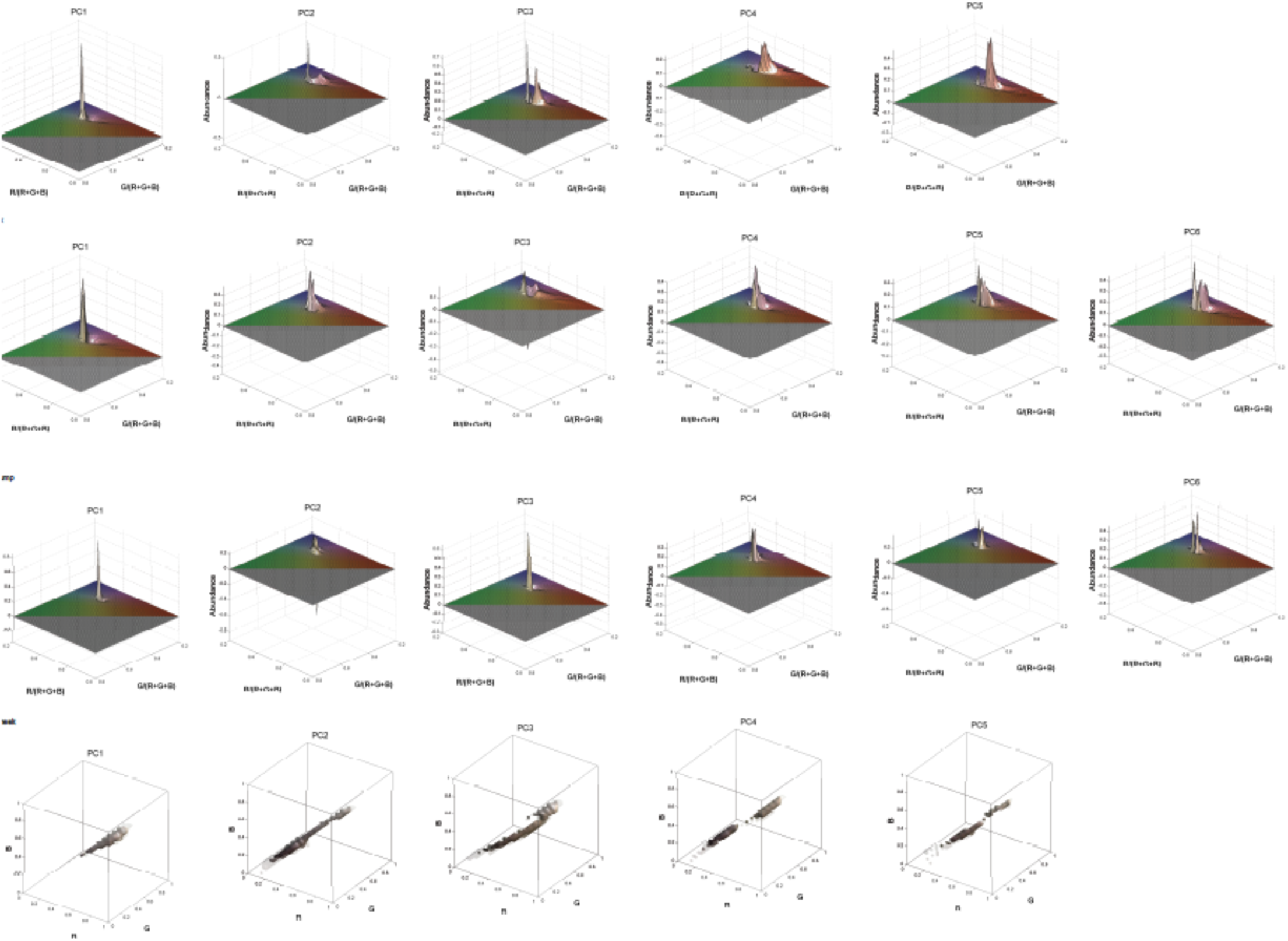
Illustration of successive eigenplanes for crown, back and rump as well as for eigenfields for cheek.

**Supplementary Figure 3.**
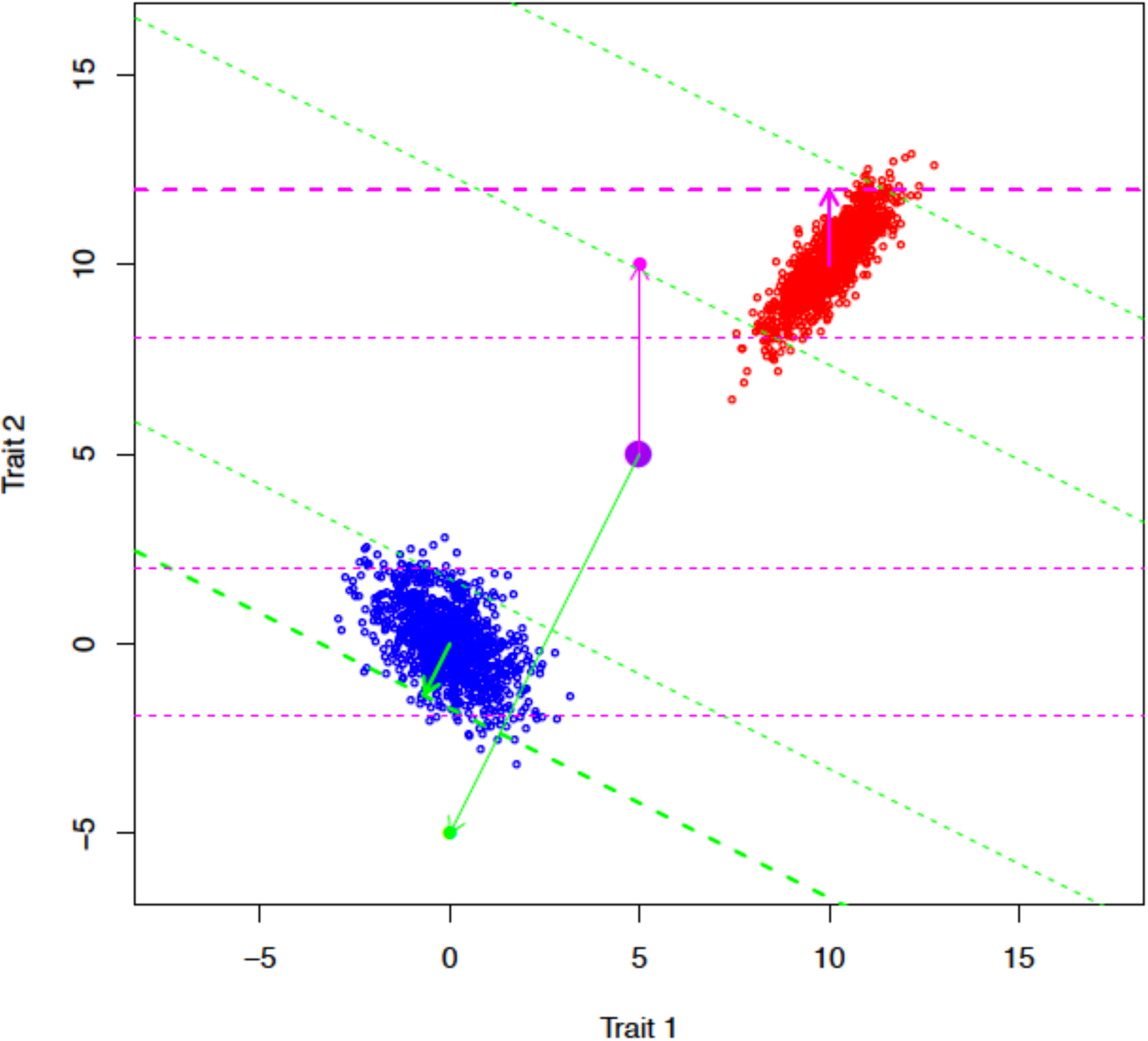
Illustration of the multivariate novelty index in the simple case of two traits. Blue and red points represent individual values for two traits for the two parental species. The large purple point is the predicted F2 mean. The green and magenta points represent trait values for two hybrid individuals. The thin green and magenta arrows beginning at the F2 mean indicate their respective axes of divergence from the predicted F2 mean; the axes used to estimate novelty. The perpendicular dashed lines show the upper and lower quantiles for the two parental species for each hybrid individual’s multivariate axis. The thick arrows commencing at parental means indicate the distance to the outer quantile of the parent species towards which the hybrid individual has evolved, and the two thick dashed perpendicular lines highlight the novelty threshold along that axis. The green individual has a novel phenotype, having a more extreme trait value than the thick green dashed line, but the magenta individual does not.

**Supplementary Figure 4.**
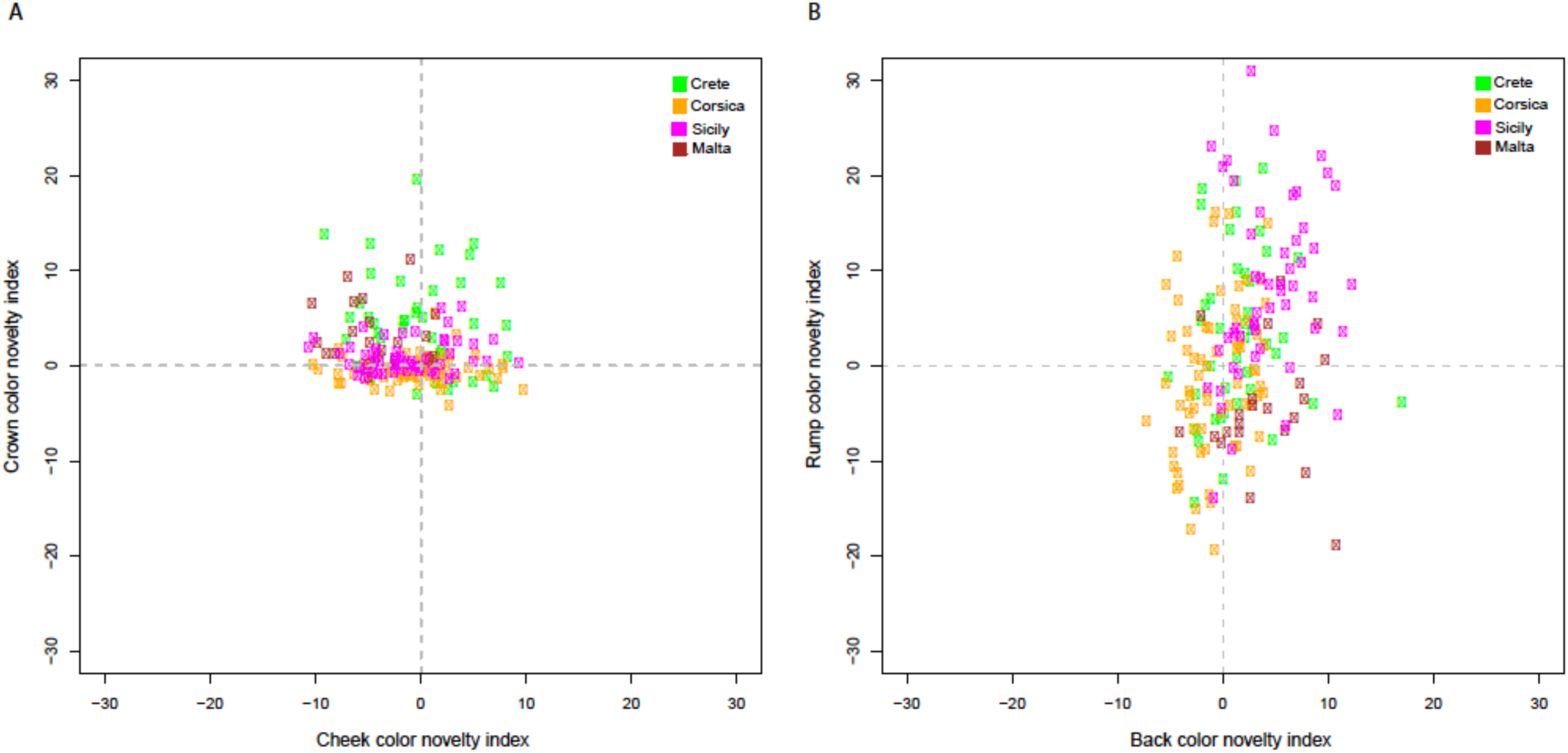
Novelty scores in Italian sparrows along the four trait axes. Each island has a different color, with green symbols representing Crete, yellow Corsica, pink Sicily and dark red Malta. Values > 0 represent novel phenotypes.

**Supplementary Table 1.**
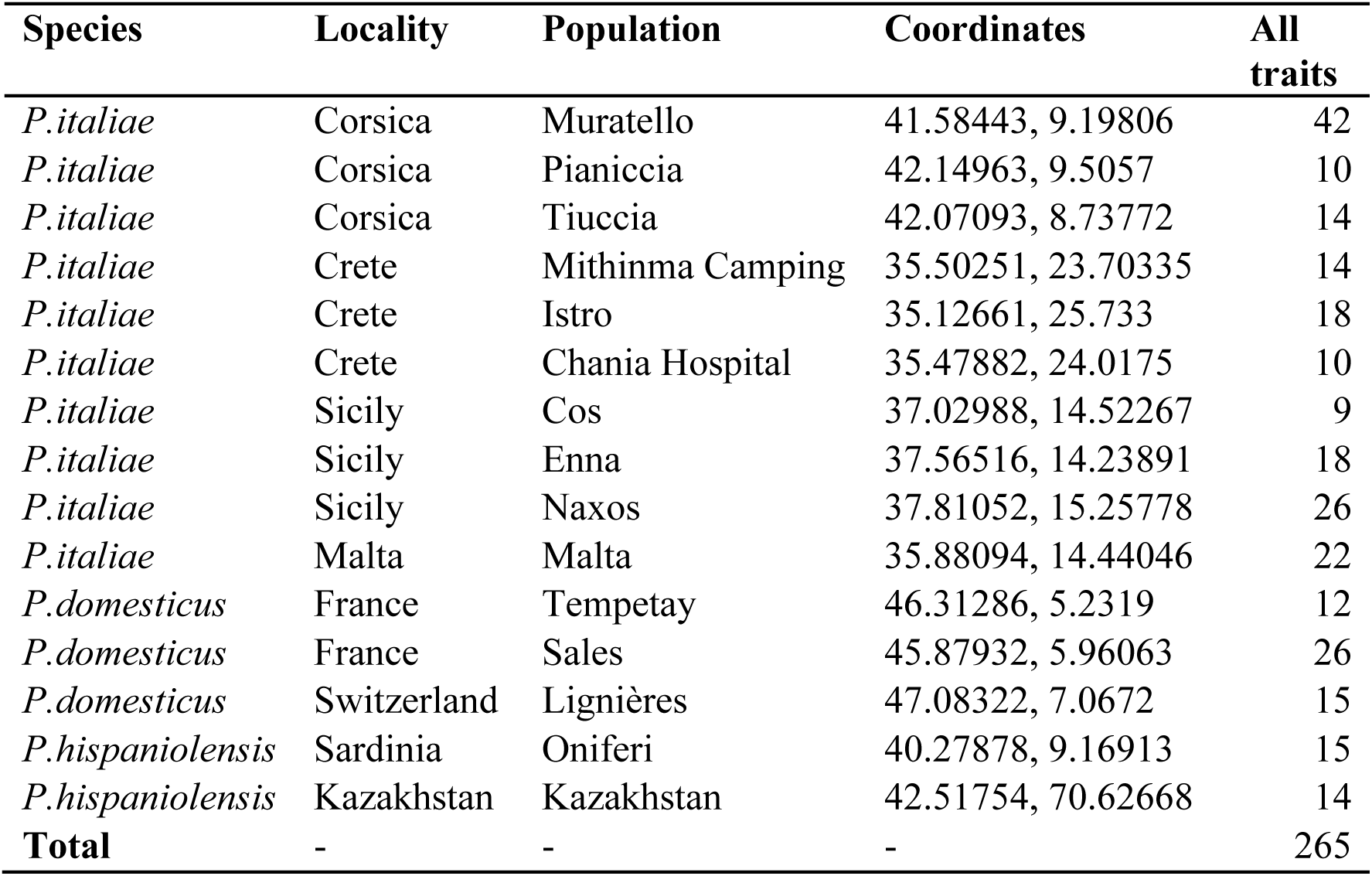
Number of individuals sampled per species, island or mainland locality and population, and coordinates in decimal degrees for the sampling locations. “All traits” means that we have sampled information for all phenotypic traits for that number of individuals.

**Supplementary Table 2.**
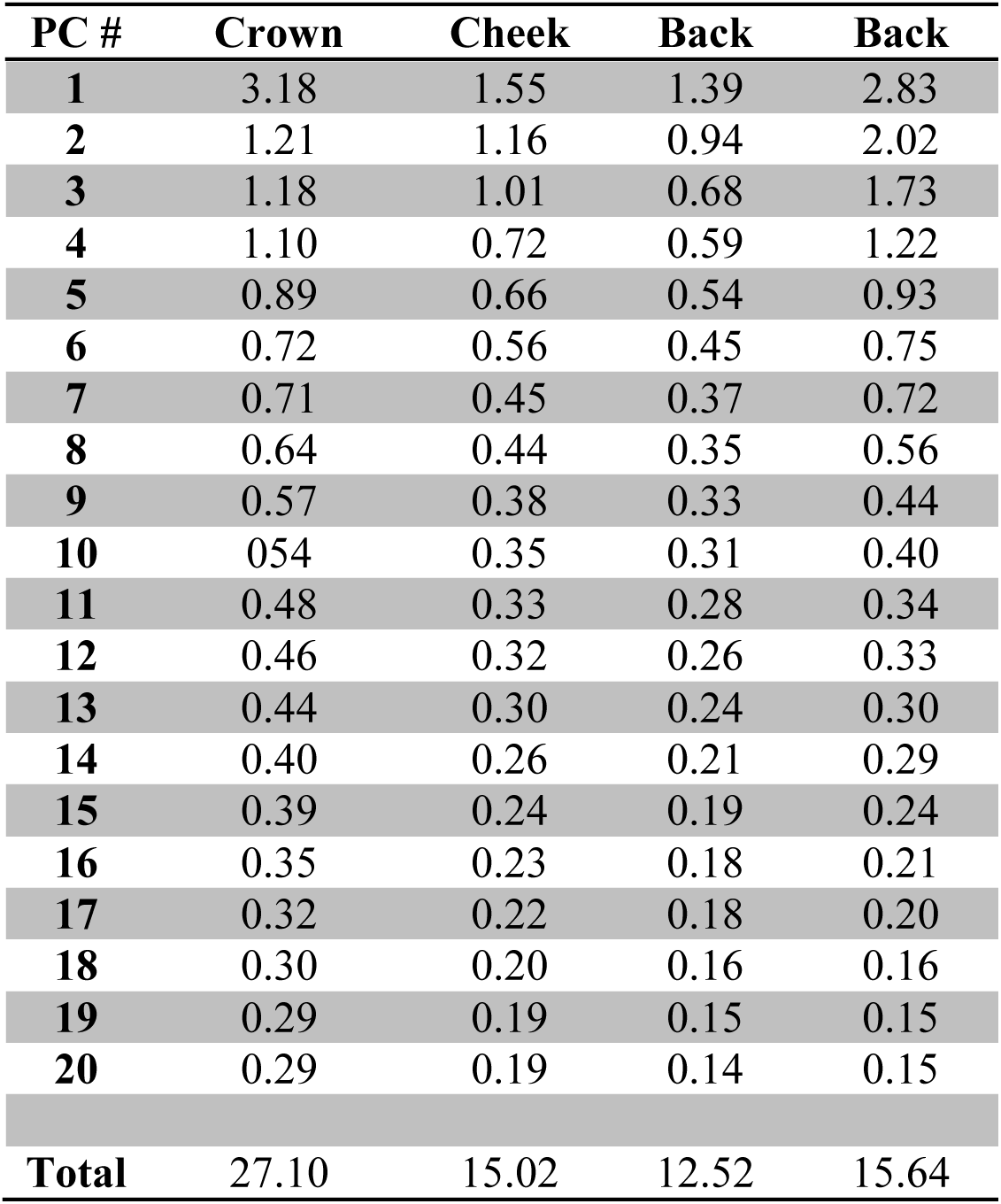
Eigenvalues for successive principal components (PCs) per study trait. The first 20 PCs are shown as the drop in eigenvalue is slow after this. The sum of the eigenvalues for all PCs is given in the Total column at the bottom.

**Supplementary Table 3.** The proportion of among-parent variance explained by within-parent evolvability. Results from a regression of among-parent variance on average within-parent evolvability along 1000 randomly chosen unit vectors.

| *Trait* | *Intercept* | *P<sub>intercept</sub>* | *Slope* | *P<sub>slope</sub>* | *R<sup>2</sup>* |
| --- | --- | --- | --- | --- | --- |
| *Crown* | 256.6 | $8 \times 10^{-16}$ | 3.5 | $3 \times 10^{-10}$ | 0.04 |
| *Back* | 39.0 | $2 \times 10^{-16}$ | -0.3 | 0.1 | 0.003 |
| *Rump* | 30.4 | 0.02 | 3.4 | $2 \times 10^{-16}$ | 0.20 |
| *Cheek* | 16.9 | $10 \times 10^{-5}$ | 0.7 | $8 \times 10^{-7}$ | 0.02 |

**Supplementary Table 4.**
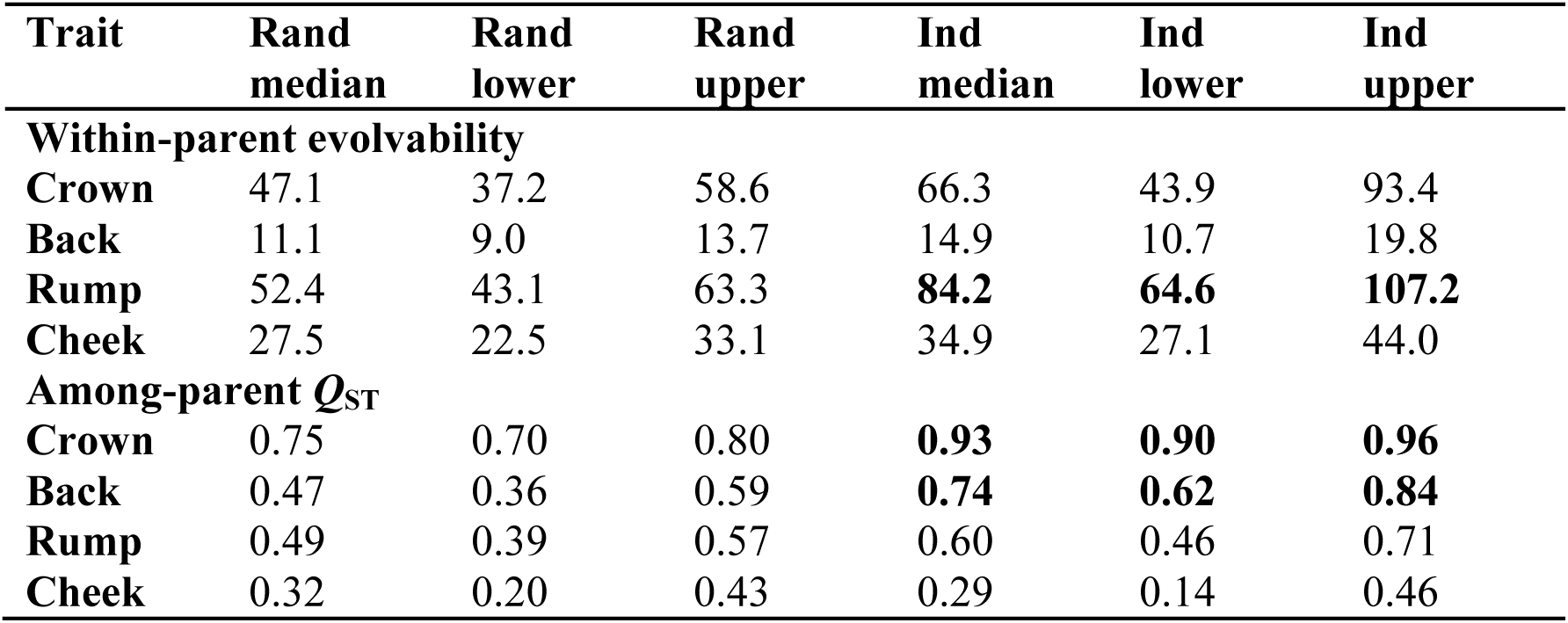
*Q*_ST_ and average within-parent evolvability for the axis of divergence of each individual from the expected F2 means and for 1000 random axes. If the 95% Highest Posterior Densities (HPD; here the lower and upper 95% are denoted lower and upper) intervals between the random and individual data set do not overlap, the difference is significant. Significant results are in bold.

| Trait | Rand<br>median | Rand<br>lower | Rand<br>upper | Ind<br>median | Ind<br>lower | Ind<br>upper |
| --- | --- | --- | --- | --- | --- | --- |
| **Within-parent evolvability** |  |  |  |  |  |  |
| **Crown** | 47.1 | 37.2 | 58.6 | 66.3 | 43.9 | 93.4 |
| **Back** | 11.1 | 9.0 | 13.7 | 14.9 | 10.7 | 19.8 |
| **Rump** | 52.4 | 43.1 | 63.3 | **84.2** | **64.6** | **107.2** |
| **Cheek** | 27.5 | 22.5 | 33.1 | 34.9 | 27.1 | 44.0 |
| **Among-parent $Q_{ST}$** |  |  |  |  |  |  |
| **Crown** | 0.75 | 0.70 | 0.80 | **0.93** | **0.90** | **0.96** |
| **Back** | 0.47 | 0.36 | 0.59 | **0.74** | **0.62** | **0.84** |
| **Rump** | 0.49 | 0.39 | 0.57 | 0.60 | 0.46 | 0.71 |
| **Cheek** | 0.32 | 0.20 | 0.43 | 0.29 | 0.14 | 0.46 |

**Supplementary Table 5.**
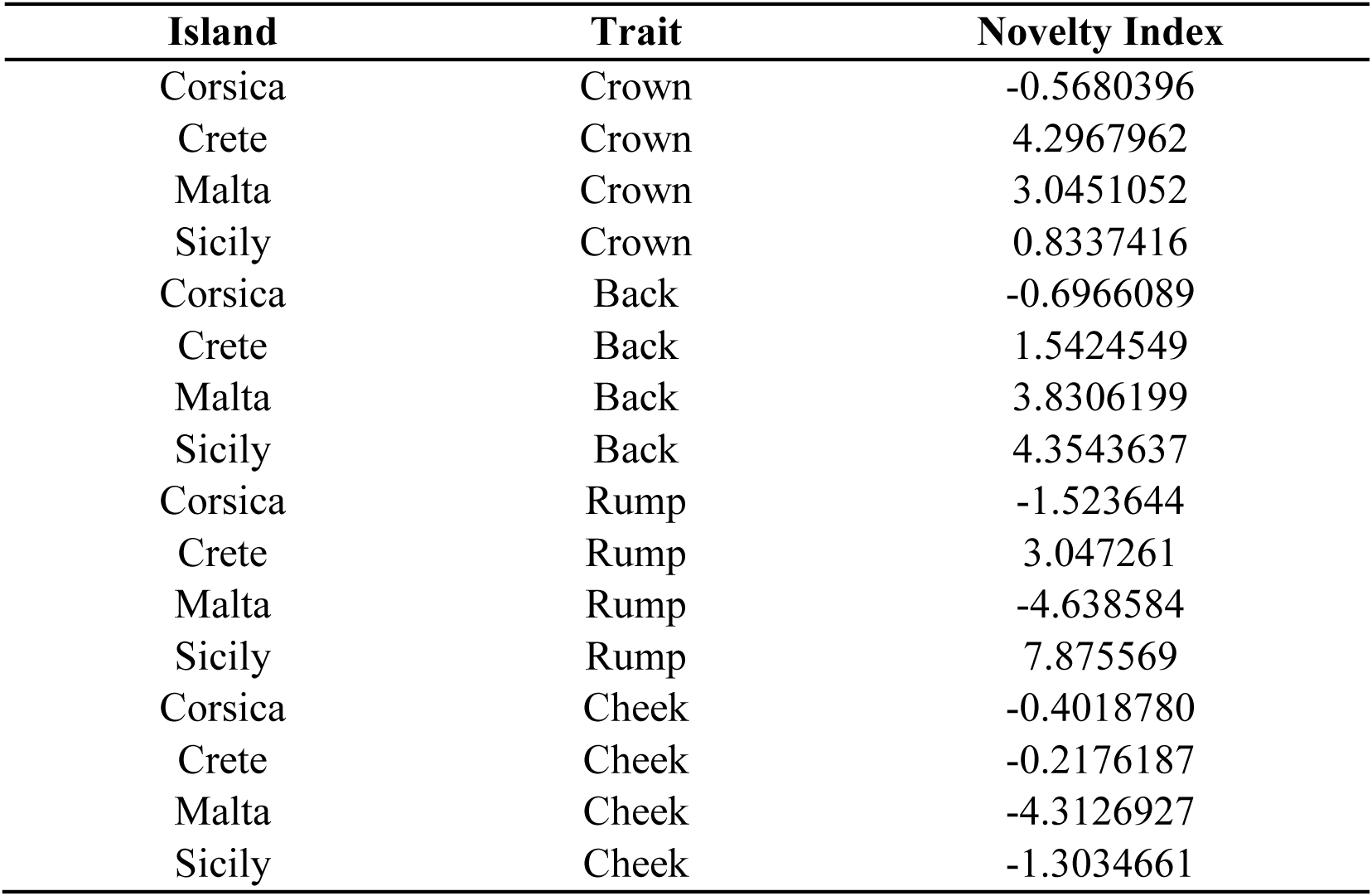
Novelty index per island population and study trait (average across individuals).

| **Island** | **Trait** | **Novelty Index** |
| --- | --- | --- |
| Corsica | Crown | -0.5680396 |
| Crete | Crown | 4.2967962 |
| Malta | Crown | 3.0451052 |
| Sicily | Crown | 0.8337416 |
| Corsica | Back | -0.6966089 |
| Crete | Back | 1.5424549 |
| Malta | Back | 3.8306199 |
| Sicily | Back | 4.3543637 |
| Corsica | Rump | -1.523644 |
| Crete | Rump | 3.047261 |
| Malta | Rump | -4.638584 |
| Sicily | Rump | 7.875569 |
| Corsica | Cheek | -0.4018780 |
| Crete | Cheek | -0.2176187 |
| Malta | Cheek | -4.3126927 |
| Sicily | Cheek | -1.3034661 |

**Supplementary Table 6.** **Multiple regression results on** the relationship between novelty index (response) and among parent *Q*_ST_ and within-parent evolvability along the axis of individual divergence from the predicted F2 phenotype.

| **Trait** | **Predictor** | **Estimate** | **Df,**<br>**error**<br>**df** | ***F*** | ***P*** | ***R*<sup>2</sup>** |
| --- | --- | --- | --- | --- | --- | --- |
| **Crown** | Among-parent $Q_{ST}$ | -0.35 | 2, 180 | 827.5 | <2.2e-16 | 0.90 |
| **Crown** | Average within-parent evolvability | -213.8 | 2, 180 |  |  |  |
| **Back** | Among-parent $Q_{ST}$ | -0.22 | 2, 180 | 13.7 | 2.90e-06 | 0.13 |
| **Back** | Average within-parent evolvability | -4.41 | 2, 180 |  |  |  |
| **Rump** | Among-parent $Q_{ST}$ | -0.023 | 2, 180 | 17.3 | 1.34e-07 | 0.16 |
| **Rump** | Average within-parent evolvability | -15.66 | 2, 180 |  |  |  |
| **Cheek** | Among-parent $Q_{ST}$ | -0.197 | 2, 180 | 20.95 | 6.60e-09 | 0.19 |
| **Cheek** | Average within-parent evolvability | -6.89 | 2, 180 |  |  |  |

**Supplementary Table 7.** Mahalanobis distances between species and their significances for all traits, as well as for the individual traits.

| **Trait** | **House-Spanish** | **House-Italian** | **Spanish-Italian** | **P(House-Spanish)** | **P(House-Italian)** | **P(Spanish-Italian)** |
| --- | --- | --- | --- | --- | --- | --- |
| **All traits** | 9.99 | 9.28 | 3.46 | 0.0003 | 0.0003 | 0.0003 |
| **Crown** | 8.63 | 8.60 | 0.97 | 0.0003 | 0.0003 | 0.7104 |
| **Back** | 2.83 | 1.09 | 2.65 | 0.0003 | 0.0003 | 0.0003 |
| **Rump** | 2.96 | 2.27 | 1.07 | 0.0003 | 0.0003 | 0.0042 |
| **Cheek** | 2.40 | 1.77 | 0.77 | 0.0003 | 0.0003 | 0.0672 |

**Supplementary Table 8.**
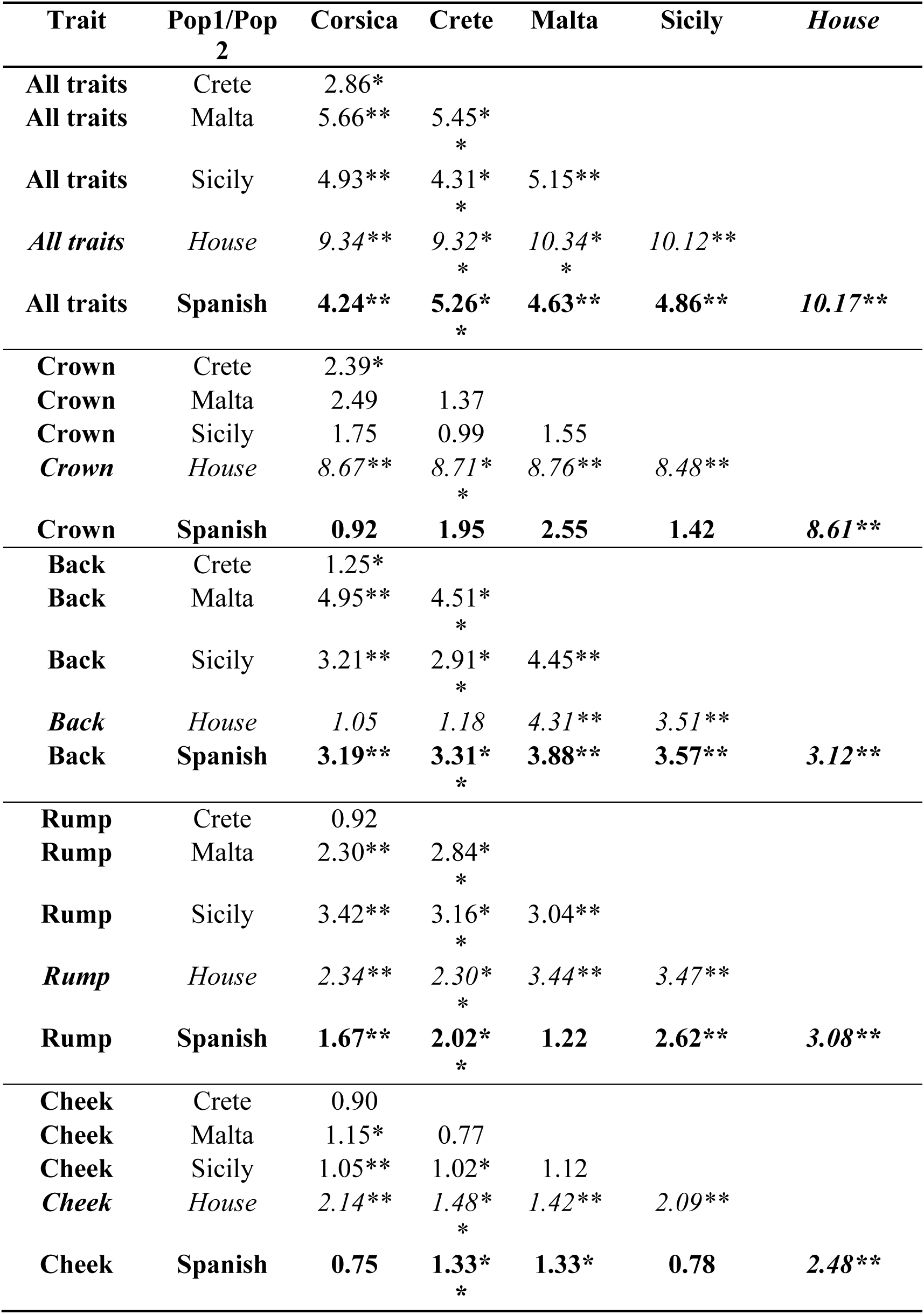
Mahalanobis distances between species and island populations of Italian sparrow and their significances for all traits, as well as for individual traits. Combinations involving house sparrow are italiziced, and these involving Spanish sparrow marked in bold. Significant divergence is denoted by stars, one star P<0.05, two stars P<0.01, three stars P<0.001.

| Trait | Pop1/Pop<br>2 | Corsica | Crete | Malta | Sicily | *House* |
| --- | --- | --- | --- | --- | --- | --- |
| **All traits** | Crete | 2.86* |  |  |  |  |
| **All traits** | Malta | 5.66** | 5.45*<br>* |  |  |  |
| **All traits** | Sicily | 4.93** | 4.31*<br>* | 5.15** |  |  |
| *All traits* | *House* | *9.34\*\** | *9.32\**<br>*\** | *10.34\**<br>*\** | *10.12\*\** |  |
| **All traits** | **Spanish** | **4.24\*\*** | **5.26\***<br>**\*** | **4.63\*\*** | **4.86\*\*** | **10.17\*\*** |
| **Crown** | Crete | 2.39* |  |  |  |  |
| **Crown** | Malta | 2.49 | 1.37 |  |  |  |
| **Crown** | Sicily | 1.75 | 0.99 | 1.55 |  |  |
| *Crown* | *House* | *8.67\*\** | *8.71\**<br>*\** | *8.76\*\** | *8.48\*\** |  |
| **Crown** | **Spanish** | **0.92** | **1.95** | **2.55** | **1.42** | **8.61\*\*** |
| **Back** | Crete | 1.25* |  |  |  |  |
| **Back** | Malta | 4.95** | 4.51*<br>* |  |  |  |
| **Back** | Sicily | 3.21** | 2.91*<br>* | 4.45** |  |  |
| *Back* | *House* | *1.05* | *1.18* | *4.31\*\** | *3.51\*\** |  |
| **Back** | **Spanish** | **3.19\*\*** | **3.31\***<br>**\*** | **3.88\*\*** | **3.57\*\*** | **3.12\*\*** |
| **Rump** | Crete | 0.92 |  |  |  |  |
| **Rump** | Malta | 2.30** | 2.84*<br>* |  |  |  |
| **Rump** | Sicily | 3.42** | 3.16*<br>* | 3.04** |  |  |
| *Rump* | *House* | *2.34\*\** | *2.30\**<br>*\** | *3.44\*\** | *3.47\*\** |  |
| **Rump** | **Spanish** | **1.67\*\*** | **2.02\***<br>**\*** | **1.22** | **2.62\*\*** | **3.08\*\*** |
| **Cheek** | Crete | 0.90 |  |  |  |  |
| **Cheek** | Malta | 1.15* | 0.77 |  |  |  |
| **Cheek** | Sicily | 1.05** | 1.02* | 1.12 |  |  |
| *Cheek* | *House* | *2.14\*\** | *1.48\**<br>*\** | *1.42\*\** | *2.09\*\** |  |
| **Cheek** | **Spanish** | **0.75** | **1.33\***<br>**\*** | **1.33\*** | **0.78** | **2.48\*\*** |

**Supplementary Table 9.** MANOVA tests of divergence between populations and species. One MANOVA was used to investigate whether the house, Spanish and Italian sparrow were differentiated from eachother, and one to test whether each of the islands were differentiated from either or both parent species or other islands.

| **Group** | **Trait** | **Df** | **F** | **Df** | **Error Df** | **P_value** |
| --- | --- | --- | --- | --- | --- | --- |
| **Species** | All traits | 2,261 | 29.53 | 44 | 482 | 1.11E-109 |
|  | Crown |  | 51.92 | 10 | 516 | 1.19E-71 |
|  | Back |  | 16.69 | 12 | 514 | 3.32E-30 |
|  | Rump |  | 16.20 | 12 | 514 | 2.59E-29 |
|  | Cheek |  | 12.48 | 10 | 516 | 1.56E-19 |
| **Islands** | All traits | 3,179 | 15.28 | 66 | 480 | 2.01E-82 |
|  | Crown |  | 11.73 | 15 | 531 | 4.22E-25 |
|  | Back |  | 27.49 | 18 | 528 | 3.24E-64 |
|  | Rump |  | 15.91 | 18 | 528 | 3.53E-39 |
|  | Cheek |  | 4.30 | 15 | 531 | 1.32E-07 |

**Supplementary Table 10.**
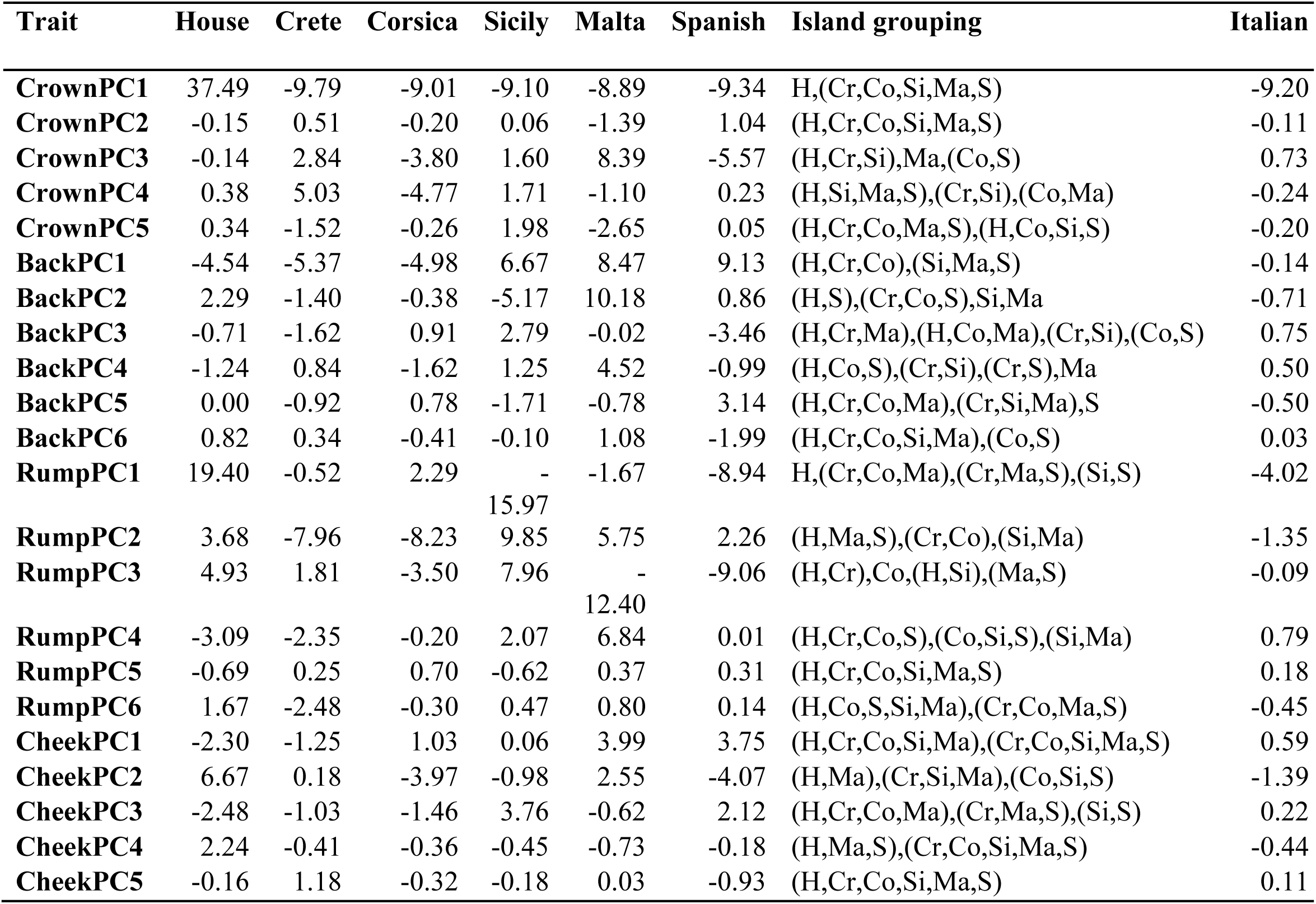
ANOVA post hoc tests of grouping between populations and species as follow-up of the MANOVAs. Parentheses denote significant differences.

**Supplementary Table 11.** Average divergence ratio across 1 000 random axes of among-island to among-population *Q*_ST_ (middle column) and average among-population *Q*_ST_ (right column).

| **Trait** | **Among vs. within island $Q_{ST}$ ratio**<br>**$\pm 95\%$ HPD:** | **Among-population $Q_{ST}$**<br>**$\pm 95\%$ HPD** |
| --- | --- | --- |
| **crown** | 3.07 (1.74, 5.08) | 0.03 (0.01, 0.04) |
| **back** | 3.66 (2.41, 5.00) | 0.10 (0.07, 0.14) |
| **rump** | 2.63 (1.75, 3.64) | 0.10 (0.06, 0.14) |
| **cheek** | 0.76 (0.36, 1.20) | 0.10 (0.06, 0.14) |

**Supplementary Table 12.** Median discriminant axis F2 quantile across all populations (middle column) and average among-island *Q*_ST_ (right column).

| **Trait** | **Median discriminant axis**<br>**F2 quantile $\pm 95\%$ HPD** | **Among island $Q_{ST}$** |
| --- | --- | --- |
| **crown** | 0.98 (0.92, 1.00) | 0.09 (0.06, 0.12) |
| **back** | 0.27 (0.17, 0.37) | 0.39 (0.32, 0.45) |
| **rump** | 0.63 (0.53, 0.74) | 0.25 (0.20, 0.31) |
| **cheek** | 0.63 (0.45, 0.79) | 0.07 (0.03, 0.11) |

## SUPPORTING METHODS

### A) Decomposition of high-dimensional color data

When there is no evidence for UV coloration, as in sparrows (Tesaker 2014), a widely used measure of coloration is the amount of Red, Green and Blue (RGB) in the pixels of an image (Gerald et al. 2001). However, mean and standard deviation of these do not take spatial correlations between colors into account (Brydegaard et al. 2012). We quantified color using the Chromatic Spatial Variance Toolbox (Brydegaard et al. 2012); available at http://www.models.life.ku.dk/ChromatricSpatialVarianceToolbox) in MatLabR2013b (version8.2.0.701; http://se.mathworks.com/products/matlab/). This method accounts for variance in spatial chromatic distributions; that color may change across a surface by patterns or patchiness. The toolbox uses the X-rite color checker in the image to standardize the coloration of each image to a uniform level, such that errors from slight illumination variation between photographs will not be included in the analysis. It then normalizes the RGB data from the standard 0-255 scale (which is based on an adjustment to reflect human visual perception) to a 0-1 scale, proportional to light reflectance (0 = no reflectance, 1 = complete reflectance).

We then use a Singular Value Decomposition (SVD) implemented in the package to decompose data. SVD2D uses the proportion of red and of green for each pixel to create a 2D histogram of color variation (Supplementary Figure 1), and therefore measures chromatic color variation while removing variation in reflectance. SVD2D performed best in correctly categorizing Skyros wall lizards (*Podarcis gaigeae*) into distinct groups based on color patterns, compared to the two other dimensionalities of SVD (Brydegaard et al. 2012), and was therefore applied to crown, rump and back yielding eigenplanes (Supplementary Figure 2). Variation in the cheek of the sparrow species is dominated by variation in reflectance intensity (white-gray-black), which is accounted for by a reflectance intensity parameter in SVD3D, and we hence applied SVD3D to cheek data. This decomposition incorporates both chromatic and reflectance variation in a 3D histogram of color variation. Only eigenvectors/planes which deviate from the noise floor are used in further analysis, and all SVD scores were mean-centered prior to subsequent analysis. After mean centering, SVD scores are equivalent to Principal Components.

### B) Evidence for directional and divergent selection

Persistent and uniform directional selection should reduce levels of divergence among populations, due to reduced polymorphism. It should also lead to extreme divergence of population means from the F2 prediction along the among-parent discriminant axis, if selection were favoring the phenotype of one parent over the other. Divergent selection should increase among-population divergence over drift expectations. As multivariate population divergence by drift scales to the G matrix (Martin et al. 2008), *Q*_ST_ should change systematically with changing within-population variance. Given the assumption that parental species are close to their phenotypic optima, we expected stronger directional or divergent selection along the among-parent discriminant axis than on average along 1000 random axes.

### C) Rationale for testing for mosaicism

If drift were a major factor in determining plumage trait values, we would expect the amount of divergence back towards the two parental phenotypes to be random among traits, and phenotypes for each trait typically to be intermediate rather than parent-like. Greater consistency across traits in evolution towards one or other parent species could be caused by among-trait genetic correlations, recent or ongoing backcrossing between hybrid and parent, or consistent selection across traits. Selection back towards different parental phenotypes in different plumage traits would produce a more mosaic phenotype than neutral expectations. Therefore, in order to strengthen our conclusion that selection is involved in hybrid trait evolution, we examined the extent of trait mosaicism: different traits having evolved back towards different parent species in the same population.

### D) Rationale for studying within-population evolvability

While average evolvability may initially be higher for all traits that are differentiated between parent species, we predict that subsequently, uniform and persistent directional selection will push hybrid evolvability as well as phenotypes of the focal Italian sparrow population, towards the values of one parent species. Whether higher average evolvability in hybrids than parents is maintained over extended periods under drift or other forms of selection is harder to predict. It depends on whether the hybrids will tend to evolve back to the same levels of polymorphism produced by mutation-selection or mutation-drift balance in the parents. However, we may predict that this process of evolution of evolvability back towards one or both parent species should be slower (and hence current hybrid evolvability values relatively higher) for traits evolving by drift than those under a strong influence of selection.

